# Diacylglycerol kinase-ε is required for the formation of GPI-anchored CD14 and modulates the LPS-induced proinflammatory responses of macrophages

**DOI:** 10.1101/2025.03.14.643240

**Authors:** Aneta Hromada-Judycka, Gabriela Traczyk, Ichrak Ben Amor, Anna Ciesielska, Katarzyna Kwiatkowska

**Affiliations:** Laboratory of Molecular Membrane Biology, Nencki Institute of Experimental Biology PAS, 3 Pasteur St., 02-093 Warsaw, Poland

## Abstract

Diacylglycerol kinase-ε (DGKε) is a unique member of the DGK family with strict specificity toward DAG containing stearic and arachidonic fatty acid residues, called SAG, and producing phosphatidic acid used for the synthesis of phosphatidylinositol (PI). PI and its derivatives, both phosphorylated and non-phosphorylated ones, regulate a multitude of processes, including the signaling of diverse plasma membrane receptors. To the latter belong Toll-like receptor 4 (TLR4) and its accessory CD14 protein activated in macrophages by bacterial lipopolysaccharide (LPS). To assess the role of DGKε in the LPS-induced pro-inflammatory responses, we obtained Raw264 cells stably depleted of DGKε and subsequently rescued them with DGKε-Myc expressed at a level similar to the native one. As a result, SAG phosphorylation was markedly decreased and then restored in those cells, with the activity of other DGKs unaffected. The depletion of DGKε nullified the LPS-induced pro-inflammatory signaling of TLR4 dependent on CD14-mediated internalization of TLR4 and the TRIF engagement in endosomes. In contrast, the MyD88-dependent signaling pathway, for which CD14 involvement can be dispensible, was inhibited only partially. In accordance, no mature, GPI-anchored form of CD14 was produced in the DGKε-depleted cells and no CD14 was found on the cell surface. The reintroduction of DGKε restored both the abundance of GPI-CD14 and the CD14-dependent signaling of TLR4. These results indicate that the DGKε-dependent phosphorylation of SAG controls the synthesis of the pool of PI that serves for the biosynthesis of the GPI moiety of CD14. We thereby have identified DGKε as a key factor determining the sensitivity of macrophages to LPS.

## INTRODUCTION

Lipopolysaccharide (LPS, endotoxin) is a major component of the outer membrane of Gram-negative bacteria belonging to so-called pathogen-associated molecular patterns (Kawai et al., 2024). Upon infection, LPS and other PAMPs trigger acute pro-inflammatory reactions aiming to eradicate the bacteria. However, when exaggerated and dysregulated, this response can lead to potentially fatal sepsis. Also, prolonged low-grade systemic endotoxemia can occur when dysbiotic microflora increase gut permeability allowing LPS to enter the bloodstream and paving the way for obesity, diabetes, and cardiovascular disease (Cani et al., 2008, Virzi et al., 2022; Violi et al., 2023). An important role in the response to LPS is played by macrophages, whose plasma membrane is equipped with a pattern-recognition receptor - Toll-like receptor 4 (TLR4) and its accessory CD14 protein. In a typical scenario, LPS aggregates released from bacteria are first recognized by serum LPS-binding protein (LBP), which facilitates subsequent binding of LPS monomers by the hydrophobic N-terminal pocket of CD14 (Kim et al., 2005, Kelley et al., 2013). The LPS monomers are then transferred from CD14 onto the complex of TLR4 with covalently linked MD2 (Ryu et al., 2017). The TLR4/MD2 complexes dimerize and recruit a pair of adaptor proteins, TIRAP and MyD88, and the latter triggers a signaling cascade involving IRAK1/2 kinases and TRAF6 E3 ubiquitin ligase, and also MAP kinases ultimately leading to the activation of NFκB and AP-1 transcription factors (Kawai et al., 1999, Park et al., 2009, Fitzgerald et al., 2001, Mothswene et al., 2009). Eventually, pro-inflammatory cytokines are expressed, including the hallmark TNFα (Bjorkbacka et al., 2004, Meissner et al., 2013). Subsequently, CD14 governs the internalization of TLR4/MD2 and in endosomes TLR4 recruits a second pair of adaptor proteins, TRAM and TRIF, launching a signaling cascade involving TRAF3 E3 ubiquitin ligase and leading to the activation of IRF3/6 transcription factors and production of type II interferons and the chemokine RANTES (Kawai et al., 2001, Yamamoto et al., 2003, Jiang et al., 2005, Husebye et al., 2006, Kagan et al., 2008, Zanoni et al., 2011). Also, a late-phase NFκB activation occurs downstream of the TRIF-TRAF6-dependent pathway (Sato et al., 2003, Cusson-Hermance et al., 2005). The LPS-induced NFκB activation also licenses a subsequent activation and assembly of NLRP3 inflammasome followed by the release of interleukin 1β and pyroptotic cell death, currently considered to be key factors in the development of (auto)inflammatory diseases (Mangan et al., 2018).

CD14 is expressed mainly in myeloid cells such as monocytes, macrophages, and dendritic cells and is embedded in the outer leaflet of their plasma membrane via glycosylphosphatidylinositol (GPI) anchor (Haziot el al., 1998, Simmons et al., 1989). CD14 is synthesized as a precursor protein (375 amino acids in human, 366 in mouse CD14) anchored in the endoplasmic reticulum membrane via its C-terminal transmembrane fragment. The C-terminal 30 amino acids are then cleaved off and replaced by a preassembled GPI moiety in a reaction catalyzed by GPI transamidase; the nascent GPI-linked protein is subjected to the GPI anchor remodeling in the endoplasmic reticulum and transported toward the plasma membrane via the Golgi apparatus, where the GPI moiety undergoes further rearrangements determining the accumulation of the protein in sphingolipid-cholesterol nanodomains (rafts) and its transport to the plasma membrane (Fujita and Kinoshita 2012, Kinoshita 2020). Concomitantly CD14 undergoes *N*- and *O*-glycosylation giving rise to several different forms of mature CD14 (Stelter et al., 1996). In resting cells, the mature plasma-membrane-bound CD14 undergoes slow constitutive endocytosis and degradation (Tan et al., 2019), however, we have recently found that some of the internalized CD14 can also recycle back to the plasma membrane, thereby replenishing the pool of CD14 ready for LPS binding (Ciesielska et al., 2022, Matveichuk et al., 2024). In addition to the plasma-membrane-bound GPI-linked CD14, soluble forms of CD14 are found in serum, facilitating the activation of even CD14-deficient cells by LPS. The mechanisms governing the generation of soluble CD14 by myeloid and nonmyeloid cells are not well understood but they are known to include shedding from the cell surface or/and exocytosis (Bazil and Strominger 1991, Metz et al., 1994, Durieux et al., 1994, Su et al., 1999, Funda et al., 2001). The functioning of the LBP - CD14 cascade allows TLR4 to be activated by low (nanomolar) concentrations of the major form of LPS, so-called smooth LPS (Gangloff et al., 2005). However, when present in high concentrations, LPS can be transferred to TLR4 also by albumin, making CD14 dispensable for the activation of the MyD88-dependent pathway (Gioannini et al., 2002, Borzecka et al., 2013). In contrast, the participation of membrane-anchored CD14 is required for the endocytosis of TLR4/MD2, and therefore also for initiating the endosomal TRIF-dependent TLR4 signaling pathway in macrophages (Jiang et al., 2005, Zanoni et al., 2011), underscoring the signaling functions of GPI-bearing CD14. In accordance, our studies have demonstrated that upon LPS binding, CD14 induces a biphasic generation of PI(4,5)P_2_ in rafts (Plociennikowska et al., 2016). A line of studies have revealed some details on how PI(4,5)P_2_ interacts with rafts and how it and products of its hydrolysis and phosphorylation control the LPS-induced endocytosis of TLR4 (Kagan and Metzhitov 2006, Aksoy et al., 2012, Chiang et al., 2012, Dong et al., 2020, Li et al., 2023).

PI(4,5)P_2_ and other derivatives of PI are important regulatory lipids controlling, among others, signal transduction including the LPS-triggered one, vesicular trafficking, non-vesicular lipid transport, and cytoskeleton organization, which makes them of substantial interest in diverse research fields (Balla 2013, Posor and Haucke 2022). The maternal molecule, PI, is also a veratile lipid used for GPI synthesis and as a source of arachidonic acid for eicosanoid production (Houjou et al., 2007, Gil de Gomez et al., 2013). PI is synthesized *de novo* in the endoplasmic reticulum through a multistep process from glycerol-3-phosphate, and the synthesis is likely combined with fatty acid remodeling in the Lands cycle to achieve the stearic and arachidonic fatty acid composition (C38:4) typical of PI (Bozelli and Epand 2019; Blunsom and Cockcroft 2020; Lee et al., 2012, Traynor-Kaplan et al., 2017, Kim et al., 2022). Additionally, PI can be generated in the so-called PI cycle taking place at the endoplasmic reticulum and plasma membrane, which can intervene with the *de novo* PI synthesis and serves to rebuild the PI level after stimulus-induced hydrolysis of PI(4,5)P_2_ into diacylglycerol (DAG) and inositol 1,4,5-trisphosphate (IP_3_) (Bozelli and Epand 2019; Blunsom and Cockcroft 2020). Despite the evident importance of PI and its derivatives for LPS-induced responses, the molecular mechanisms controlling their abundance and turnover, and their relation to CD14 are far from being elucidated.

Studying those questions, we have found that several enzymes involved in the PI cycle are palmitoylated in an LPS-dependent manner (Sobocinska et al., 2018). These include diacylglycerol kinase-ε (DGKε) whose palmitoylation was previously unknown (Sobocinska et al., 2018). DGKε is a member of a family of ten enzymes phosphorylating DAG to phosphatidic acid (PA), and thereby controlling the cellular level of these two key intermediates in lipid synthesis, with some of their species also functioning as signaling molecules (Sakane et al., 2024). Among all the DGKs only DGKε is an integral membrane protein anchored in the membrane via an N-terminal α-helix (Traczyk et al., 2024). We found that DGKε is *S*-palmitoylated at Cys 38/40 (mouse/human) of this α-helix (Traczyk et al., 2024). Importantly, only DGKε displays strict specificity toward DAG bearing the C18:0 (stearic) and C20:4 (arachidonic) fatty acids at the *sn*-1 and *sn*-2 position, respectively, and therefore abbreviated SAG, giving rise to 1-stearoyl-2-arachidonoyl-phosphatidic acid (SAPA) (Tang et al., 1998, Pettitt and Wakelman 1999, Lung et al., 2009, Ware et al., 2020). As the predominant form of PI bears exactly the same C38:4 fatty acids, it can be derived in the PI cycle from SAPA which in turn is synthesized from SAG, suggesting a critical role of DGKε in an unhindered operation of the PI cycle (Epand et al., 2016). However, a contribution of DGKs other than DGKε to the PI cycle has been implicated recently, with a concomitant suggestion that the cycle also provides PI in resting cells (Kim et al., 2022, Bernada et al., 2022). To examine the contribution of DGKε to LPS-induced signaling, we obtained Raw264 cells depleted of DGKε by shRNA silencing and subsequently rescued them with Myc-tagged DGKε. Unexpectedly, we found that DGKε is indispensable for the formation of mature, GPI-anchored CD14, likely by controlling the synthesis of the GPI anchor. Consequently, DGKε turned out to affect the TRIF-dependent signaling of TLR4 and also, to a lower extent, the MyD88-dependent one. We thereby identified DGKε as a key factor affecting the response of macrophages to LPS.

## MATERIALS AND METHODS

### Cell culture and stimulation

Raw264.7 (further Raw264) cells (ATCC) were cultured in DMEM containing 10% FBS (Thermo Fisher Scientific) and 4.5 g/l glucose. Cells were stimulated with 10 or 100 ng/ml smooth LPS from *Escherichia coli* O111:B4 (List Biological Laboratories) at 5% CO_2_, 37°C for up to 4 h.

### Construction of Raw264 cells with silenced and rescued expression of *Dgke*

To obtain Raw264 cells depleted of DGKε, they were transfected with lentiviral particles (at MOI = 1 or 5) containing five different sequences encoding *Dgke*-targeting shRNA (Merck, Table 1). Cells infected with transduction particles bearing non-mammalian shRNA (at MOI = 1 or 5) (Merck, Table 1) served as controls. The cells were cultured in the presence of 2 μg/ml puromycin until the mortality of non-infected cells reached 100%, leaving only cells transfected with shRNA alive, essentially as described earlier (Traczyk et al., 2022, Matveichuk et al., 2024). To reintroduce DGKε, cells transfected at MOI = 5 with shRNA No. 1 and selected as described above were transfected (at MOI = 5) with lentiviral particles bearing the DGKε cDNA sequence with a double Myc tag added at the C-terminus, and conferring resistance to G418. The cDNA sequence was designed to contain eight point mutations in six codons which did not change the amino acid sequence of DGKε but protected its mRNA from degradation guided by the previously introduced shRNA (Table 1). Cells were cultured in the presence of 0.3 mg/ml G418 and selected as in the first round of transfection to obtain DGKε-Myc-rescued cells. In parallel, cells depleted of DGKε with shRNA No. 1 and also control cells obtained after the first round of transfection and selection (MOI = 5) were subjected to a second round of transfection with lentiviral particles bearing the G418 resistance gene only (at MOI = 5) and selected to obtain the ultimate DGKε-KD and control (Ctrl) cells. Lentiviral particles bearing the pLenti-C-Myc-DGGK-IRES-Neo vector, with or without the DGK-Myc coding sequence, were custom-made by Origene. The efficiency of *Dgke* silencing and its reversion was verified with RT-qPCR using primers specific to the mouse *Dgke* gene, with *Tbp* as a reference (Table 2, see also Traczyk et al., 2022).

**Table 1.**
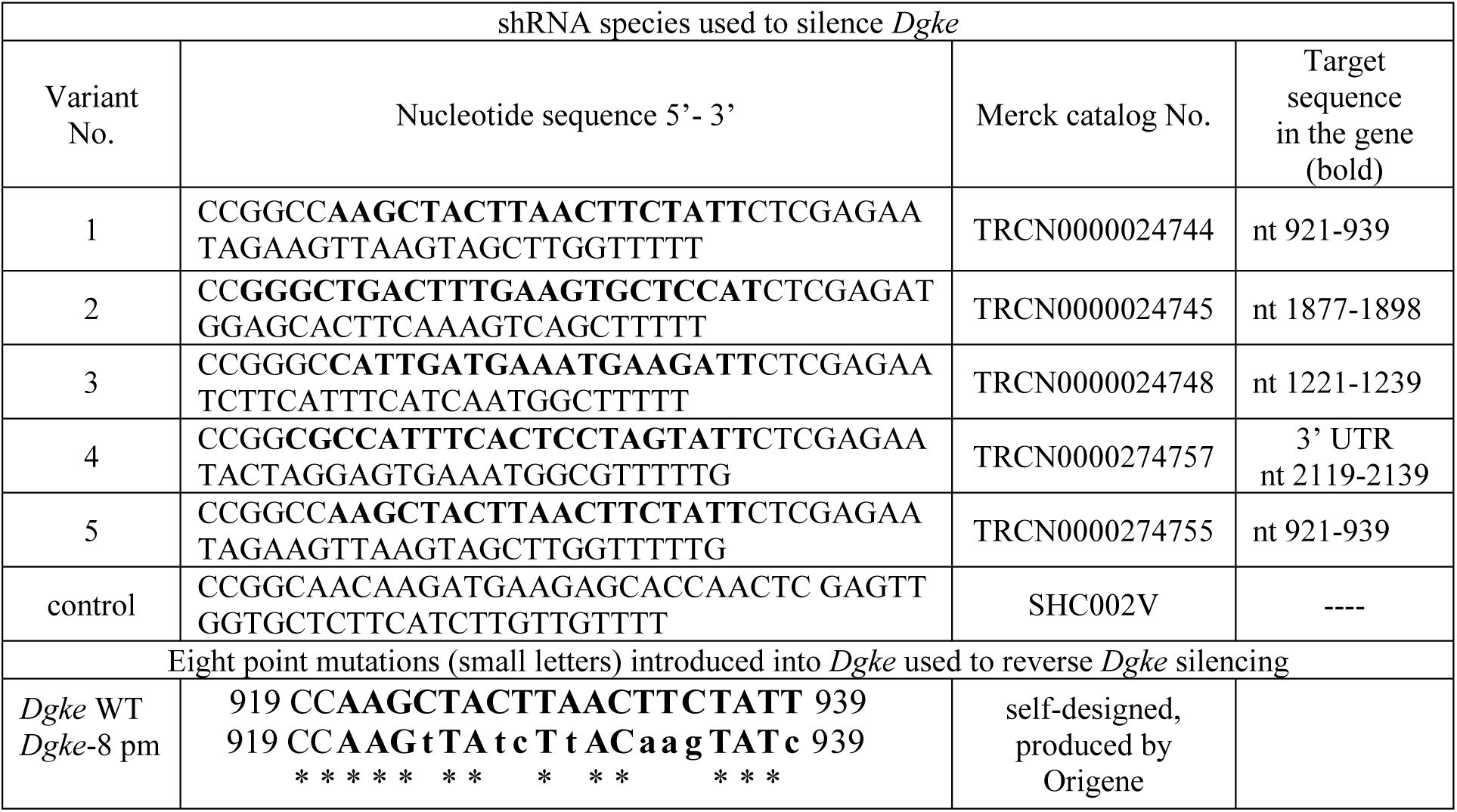
shRNA species used to silence *Dgke* and nucleotide sequence of the *Dgke* fragment bearing silent point mutations used to reverse the *Dgke* silencing.

**Table 2.**
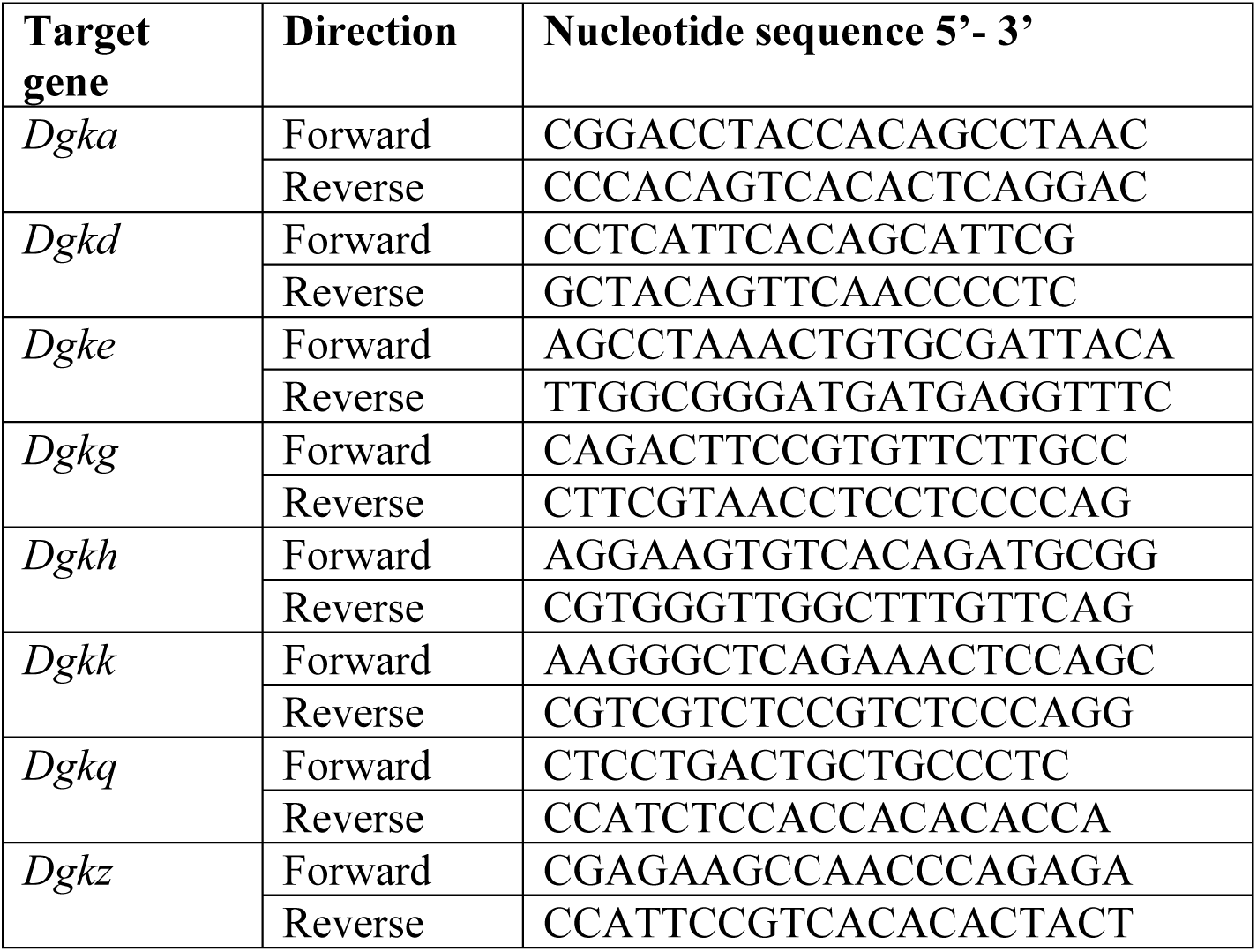
Primers used for RT-qPCR analysis of *Dgka, d, e, g, h, k, q* and *z* expression.

### Flow cytometry

Cell-surface CD14 and TLR4 were analyzed by flow cytometry using antibodies listed in Supplemental Table 1 as described in (Matveichuk et al., 2024). Total GPI-anchored proteins were stained with fluorescence-labeled inactive toxin aerolysin (FLAER) (Spark Blue 488 FLAER, Biolegend, cat. No. 567904) in 1% FBS for 30 min, according to the manufacturer’s instruction. In a series of experiments, cells were treated with 0.2 U/ml phosphatidylinositol-specific phospholipase C (PI-PLC, Invitrogen, cat. No. P6466) for 1 h at 37°C (Plociennikowska et al., 2016) before the labeling. Cell fluorescence was determined with a Becton Dickinson FACS Calibur flow cytometer. FITC/Alexa488 and phycoerythrin fluorescence was detected using a 530/30 nm and a 585/42 nm filter with an FL-1 and FL-2 detector, respectively. Data were analyzed using BD CellQuest Pro software (BD Biosciences) and the amounts of cell-surface CD14, TLR4, and GPI-anchored proteins were calculated based on the geometric mean of fluorescence intensity, as described earlier (Matveichuk et al., 2024).

### Fractionation of TX-100 lysates of cells

Cells were counted (1 × 10^6^ per sample) and fractionated into Triton X-100 (TX-100)-soluble, TX-100-insoluble (also called detergent-resistant (DRM)), and SDS-soluble fractions essentially as described earlier, with the DRM fraction corresponding roughly to membrane rafts (Matveichuk et al., 2024). Equivalent volumes of the fractions were subjected to SDS-PAGE followed by immunoblotting.

### Extraction of GPI-anchored proteins with TX-114

To analyze the partition of membrane proteins with TX-114, the detergent (Thermo Fisher Scientific cat. No 422360025) was precondesed to 12% according to Taguchi and Schatzl (2014). Cells (2 × 10^6^) were homogenized by sonication in 200 μl of the homogenization buffer (5 mM EDTA, 1 mM PMSF, 20 μg/ml aprotinin, 20 μg/ml leupeptin, 20 mM Tris, pH 7.4), supplemented with 250 mM sucrose and subjected to centrifugation (1,000 x *g*, 10 min, 4°C) to obtain the post-nuclear supernatant (PNS). Next, the PNS was ultracentrifuged (200,000 x *g*, 1 h, 4°C) and the obtained supernatant containing cytosolic proteins was collected. The pellet (membrane fraction) was dissolved in 200 μl of the PI-PLC reaction buffer containing 20 mM Tris, 0.1% TX-100, protease inhibitors as above, pH 7.5 (Fujita et al., 2022). The lysate was divided in half and supplemented or not with 2 U/ml PI-PLC. After 1 h (37°C), TX-114 was added to both samples to 2% final concentration and after a 10-min incubation on ice, the samples were centrifuged (1,000 x *g*, 10 min, 37°C) and separated into aqueous (upper) and detergent (lower) phase, the latter washed once by centrifugation with PI-PLC reaction buffer (37°C); both samples were supplemented with respective buffer to keep sample conditions and volumes consistent (Fujita et al., 2022). Proteins were precipitated from each fraction with methanol:chloroform:water (3:1:4, v:v:v), dissolved in SDS-sample buffer, and equivalent volumes were subjected to SDS-PAGE.

### DGKε activity assay

DGKε activity was determined using the mixed micelle activity assay with 1-NBD-stearoyl-2-arachidonoyl-*sn*-glycerol (NBD-SAG, Cayman Chemical, cat. No. 10011300) developed and characterized by us earlier (Traczyk et al., 2022, 2024). Briefly, cells (6 × 10^5^ per sample) were lysed in 200 μl of lysis buffer (1% NP-40, 150 mM NaCl, 1 mM EDTA, 1 mM EGTA, 1 mM DTT, 1 mM PMSF, 20 μg/ml aprotinin, 20 μg/ml leupeptin, 20 mM Tris-HCl, pH 7.4) and sonicated (Traczyk et al., 2022). Subsequently, 50 μl of the lysate containing 40 or 50 μg of total protein was used for the activity assay with 50 μl of micelle suspension, 100 μl of H_2_O and 0.5 mM ATP. The micelles were composed of NBD-SAG/SAG (at a 1:9 molar ratio, both from Cayman Chemical, cat. Nos. 10011300 and 10008650) and 1,2-dioleoyl-*sn*-glycero-phosphoserine (DOPS, Merck, cat. No. 840035C) at a final concentration of 1.45:2.03 mol% in 4 x reaction buffer containing 400 mM NaCl, 80 mM MgCl_2_, 4 mM EGTA, 4 mM DTT, 300 mM octyl-β-glucoside, 200 mM MOPS, pH 7.2. The reaction was carried out for 10 min (24°C), then lipids were extracted and separated by TLC on silica gel 60 (Merck) (1/5-1/10 of the reaction mixture) along with standard, 0.5-50 pmol 1-palmitoyl-2-NBD-dodecanoyl-*sn*-glycero-3-phosphate (NBD-PDPA, Avanti, cat. No. 810174), with chloroform:methanol:acetic acid 80% (65:15:5, v:v:v) as a mobile phase. NBD-lipids, i.e., 1-NBD-stearoyl-2-arachidonoyl-*sn-*glycero-3-phosphate (NBD-SAPA) and NBD-PDPA were visualized using G:Box (Syngene) and fluorescence intensity was assessed with ImageJ software using a standard curve drawn for NBD-PDPA and corrected by background subtraction.

In a series of experiments, DGKε activity and the activity of other DGKs were determined in cell fractions. For this purpose, cells (3 × 10^6^) were homogenized (1 mM EDTA, 1 mM EGTA, 1 mM DTT, 1 mM PMSF, 20 μg/ml aprotinin, 20 μg/ml leupeptin, 20 mM Tris-HCl, pH 7.4), supplemented with 250 mM sucrose, centrifuged to obtain PNS, which was then supplemented with 1 M NaCl and ultracentrifuged, as described above. The obtained supernatant containing cytosolic proteins was collected, pellet (membrane fraction) was dissolved in homogenization buffer supplemented with 1 M NaCl and protein concentration was determined in all fractions. Subsequently, 40 μg of total protein was used for the DGKε activity assay and analyzed, as described above for lysates. In another set of experiments, treatment of samples with 1 M NaCl was omitted. Alternatively, to analyze the activity of other DGKs, mixed micelles containing 1-NBD-decanoyl-2-decanoyl-*sn*-glycerol/1,2-didecanoyl-*sn*-glycerol (at a 1:9 molar ratio, both from Cayman Chemical, cat. Nos. 9000341 and 62210) were used and the reaction buffer was supplemented with 10 mM CaCl_2_ instead of 1 mM EGTA.

In parallel, cell lysates and fractions were supplemented with 2% SDS, vortexed and incubated at room temperature for 15 min, supplemented with 1 x SDS-sample buffer (final concentration), incubated again for 15 min, heated for 10 min at 95°C and subjected to SDS-PAGE and immunoblotting (antibodies listed in Supplemental Table 1).

### RNA isolation and RT-qPCR

RNA was isolated from cells using the Universal RNA purification kit (EURx) and reverse-transcribed into cDNA using the High Capacity cDNA Reverse Transcription Kit (Thermo Fisher Scientific) according to the manufacturer’s instructions. qPCR was performed in a StepOnePlus instrument using Fast SYBR Green Master Mix (Thermo Fisher Scientific). The primers and the PCR conditions were described earlier (Traczyk et al., 2022, Matveichuk et al., 2024) or are shown in Table 2. mRNA levels for investigated genes were calculated by the ΔΔCT method relative to the mRNA level for the *Tbp* or *Hprt* gene, each variant run in triplicate.

### Cytokine assays

TNFα and CCL5/RANTES were quantified in cell culture supernatants using mouse ELISA kits (BioLegend, R&D Systems). The profile of all secreted inflammatory markers was assayed using Mouse Cytokine Array Kit, Panel A (R&D), as described earlier (Ciesielska et al., 2021, Matveichuk et al., 2024).

### Immunoblotting

Proteins were separated by 10% SDS-PAGE and transferred to nitrocellulose membranes which were subsequently probed with antibodies listed in Supplemental Table 1. Immunoreactive bands were detected by chemiluminescence and analyzed densitometrically using the ImageJ program, as described earlier (Sobocinska et al., 2018, Prymas et al., 2020, Matveichuk et al., 2024).

### Data analysis

The significance of differences was calculated using one-way or two-way ANOVA with Tukey’s or Scheffe’s post hoc test, or Student’s *t-*test, as indicated in figure legends. For clarity, not all of the significant differences are marked in the figures.

## RESULTS

### Characteristics of macrophages depleted of DGKε and rescued with DGKε-Myc

To reveal the role of DGKε in LPS-induced signaling, we obtained Raw264 cells with stably silenced expression of *Dgke* using shRNA in lentiviral particles followed by puromycin selection. Among the five shRNA species used, variants No. 1 and No. 5 were the most effective in reducing DGKε mRNA in both resting and LPS-stimulated cells, while control non-mammalian shRNA did not interfere with the DGKε mRNA level (Fig. 1A, Supplemental Fig. 1). Subsequently, we re-introduced DGKε Myc-tagged at the C terminus to the Raw264 transfectants depleted of DGKε with shRNA variant No. 1, and selected the rescued cells by G418 resistance. In parallel, the respective control and DGKε-depleted cells were subjected to the second round of transfection with an empty vector carrying only the G418-resistance gene. In this manner we obtained: control (Ctrl) cells, in which the DGKε expression was unaffected compared to maternal Raw264 cells, DGKε knock-down cells (DGKε-KD) with the relative DGKε mRNA level reduced by about 85%, and cells expressing DGKε-Myc at a level comparable to that of endogenous DGKε (exceeding it about 2.3-fold), called the DGKε-Myc-rescued variant (Fig. 1A). We confirmed the production of DGKε-Myc protein in the rescued cells (Fig. 1B). The down-regulation of DGKε in DGKε-KD cells reduced the phosphorylation of SAG to SAPA in cell lysates by about 41-48% (*vs.* wt and Ctrl cells). It was fully restored in DGKε-Myc-rescued cells, surpassing the activity in control cells 1.5-1.7-fold (Fig. 1C-D).

**Figure 1.**
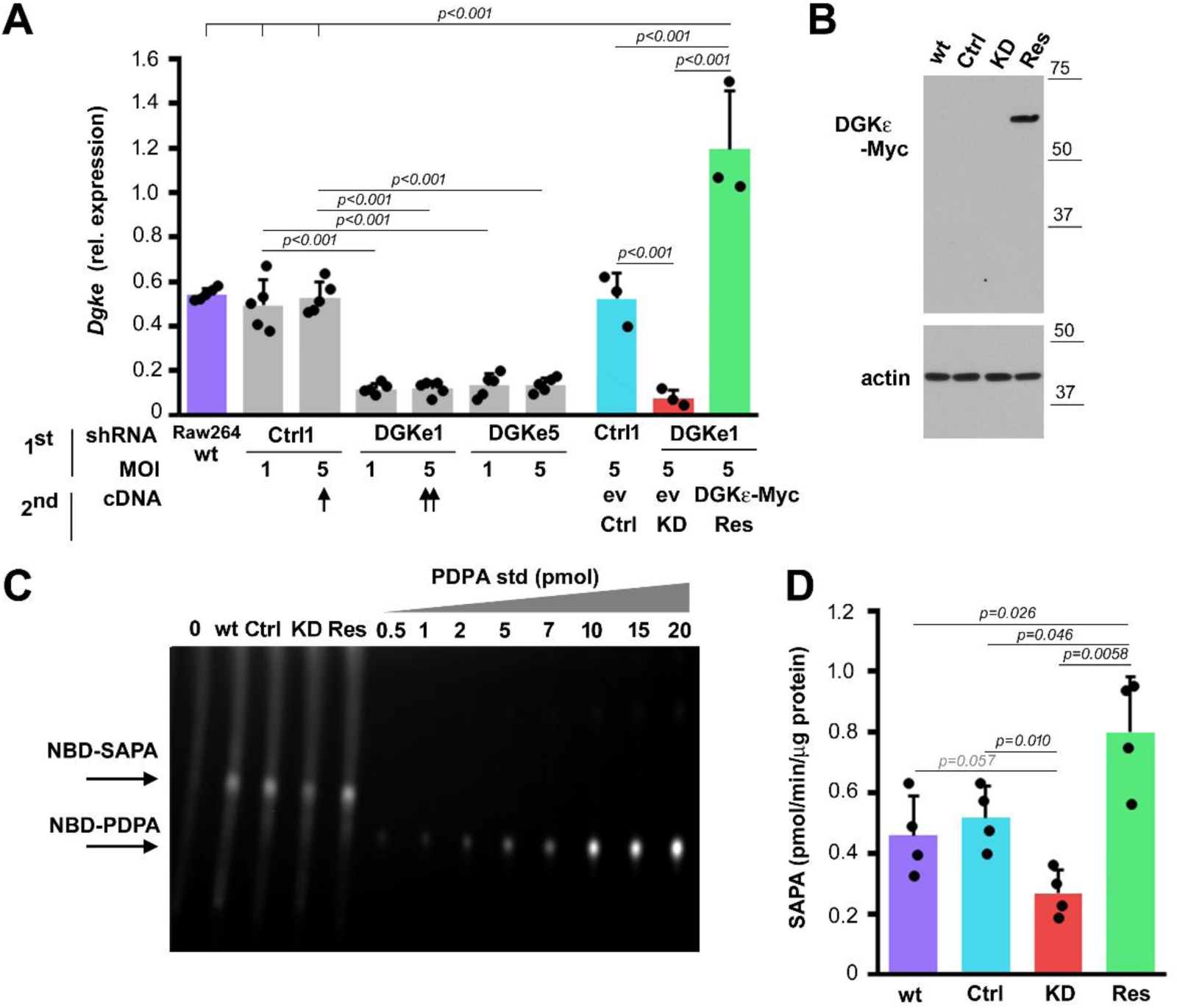
Knock-down of *Dgke* and its rescue with DGKε-Myc in Raw264 cells. (A) RT-qPCR analysis of DGKε mRNA. To silence *Dgke* expression, commercially available lentiviral particles bearing different shRNA variants were used individually (DGKe1, DGKe5) at MOI = 1 or 5 followed by puromycin selection. In parallel, control shRNA (Ctrl1) was applied at MOI = 1 or 5. Wild-type Raw264 cells – wt Raw264. In the rescue experiment, the marked DGKe1 cell variant (double arrow) was transfected with lentiviral particles bearing a sequence encoding DGKε-Myc (MOI = 5). In parallel, the Ctrl1 cells (arrow) and the DGKe1 cells were transfected with lentiviral particles bearing the antibiotic resistance gene only (ev) (MOI = 5). The three cell variants were subjected to G418 selection. Ultimately, control (Ctrl), DGKε-KD (KD) and DGKε-Myc-rescued (Res) cells were obtained. (B) Production of DGKε-Myc in DGKε-Myc-rescued cells revealed by immunoblotting with anti-Myc antibody in cell lysates. Positions of molecular weight markers are shown on the right in kDa. (C, D) Phosphorylation of SAG to SAPA in cell lysates by the fluorescence assay. (C) Representative TLC separation revealing NBD-SAPA production. The reaction mixture contained 50 μg of total protein, lipids from 1/5 of the reaction mixture were applied onto the plate. NBD-labeled lipids were separated by TLC together with 0.5–20 pmol NBD-PDPA used to draw a standard curve for each experiment. “0” – no lysate added. (D) SAPA production based on densitometric analysis of NBD-SAPA and the calibration curve for NBD-PDPA. Data shown are mean ± SD. Significantly different values as indicated by 1-way ANOVA with Scheffe’s post hoc test (A) or Student’s *t*-test (D) are marked. Additionally, in (A), wt Raw264 cells are significantly different from DGKe1, DGKe5, and DGKε-KD cells (not marked).

The SAG-specific activity remaining in DGKε-KD cell lysates can be ascribed to DGKs other than DGKε whose accessibility to SAG was facilitated by the cell lysis. Among the six *Dgk* genes likely to be expressed in monocytes/macrophages in addition to *Dgke* (Yamamoto et al., 2014), *Dgkz*, *Dgkd,* and *Dgkh* (encoding DGKη) were found to be expressed relatively abundantly in Raw264 and Ctrl cells, exceeding or being comparable to the level of *Dgke*, while *Dgka*, *Dgkg,* and *Dgkq* (encoding DGKθ) showed low expression (Fig. 2A, Supplemental Fig. 2). No expression of *Dgkκ* was found, as could be expected; *Dgkβ* and *Dgkι* are expressed specifically in neuronal cells (Yamamoto et al., 2014, Shirai and Saito, 2014). Interestingly, expression of *Dgka* and *Dgkg* was reduced in DGKε-KD cells and returned to the control level in DGKε-Myc-rescued cells (Supplemental Fig. 2). To address the issue of the possible contribution of DGKs other than DGKε to the SAG phosphorylation, we fractionated homogenates of the studied cells into cytosolic and membrane fractions in the presence of 1 M NaCl to extract DGKs possibly bound to the surface of membranes (Kai et al., 1994; Nagaya et al., 2002). Under these conditions, nearly all DGKζ was found in the cytosolic fraction marked by the presence of IκB (Fig. 2B), and DGKε was located exclusively in the membrane fraction, as indicated by the distribution of DGKε-Myc and calnexin, a transmembrane protein of the endoplasmic reticulum (Fig. 2B). Notably, the activity of the cytosolic DGKs toward SAG was not changed in DGKε-KD or DGKε-Myc-rescued cells compared with controls (Fig. 2C, upper panel; and Fig. 2D). In contrast, the activity of DGKε in the membrane fraction was lower by about 80% in DGKε-KD cells *vs.* controls and in DGKε-Myc-rescued cells its level was about 2.5-fold higher than in the controls, closely reflecting the levels of DGKε mRNA in these cells (Fig. 2C, upper panel; and Fig. 2D, compare with Fig. 1A). For comparison, the membrane fraction not treated with 1 M NaCl contained all the DGKζ, and the phosphorylation of SAG in this fraction was, on average substantially higher than in the corresponding cytosolic one, except for the membranes of DGKε-KD cells, where the its inhibition was clearly detectable (Supplemental Fig. 3A, compare with Fig. 2B). This indicated the effectiveness of 1 M NaCl in dissociating DGKs, other than DGKε, from the membrane surface and their shift to the cytosolic fraction.

**Figure 2.**
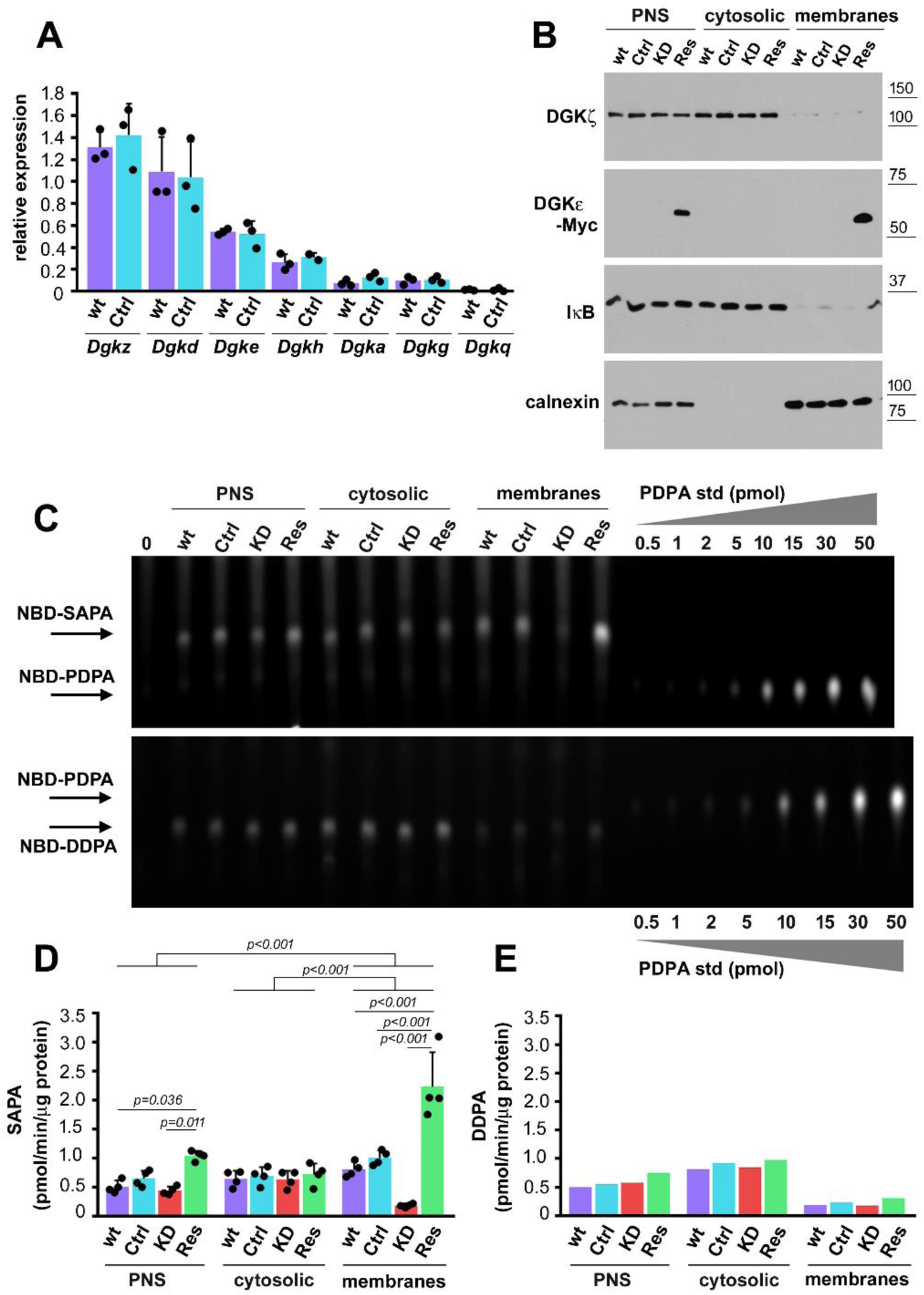
SAG phosphorylation is affected by *Dgke* knock-down and rescue. Post-nuclear supernatants (PNS) of wild-type Raw264 (wt), Ctrl, DGKε-KD, and DGKε-Myc-rescued cell homogenates were fractionated into cytosolic and membrane fractions in the presence of 1 M NaCl. (A) Expression of *Dgkz, d, e, h, a, g, and q* in Raw264 and Ctrl cells. Respective transcripts were quantified by RT-qPCR relative to *Tbp.* (B) Abundance of indicated proteins in cell fractions determined by immunoblotting. Positions of molecular weight markers are shown on the right in kDa. (C-E) Phosphorylation of SAG to SAPA (upper panel in C, D) and 1,2-didecanoyl-*sn*-glycerol (DDG) to 1,2-didecanoyl-3-phosphatidic acid (DDPA) (upper panel in C, E) in cell fractions determined using the fluorescence assay. (C) Representative TLC results revealing NBD-SAPA (upper panel) and NBD-DDPA (upper panel) production. The reaction mixture contained 40 μg of total protein, lipids from 1/5 of the reaction mixture were applied onto the plate. NBD-labeled lipids were separated by TLC together with 0.5–50 pmol NBD-PDPA used to draw a standard curve for each experiment. “0” – no homogenate added. (D, E) SAPA (D) and DDPA (E) production based on densitometric analysis of SAPA or DDPA and calibration curve for NBD-PDPA. Data shown are mean ± SD. Significantly different values as indicated by two-way ANOVA with Tukey’s post hoc test are marked.

Furthermore, in the salt-stripped membranes only a low level of phosphorylation of another DAG species, 1,2-didecanoyl-*sn*-glycerol (DDG), was detected in the membrane fraction, in line with the specificity of DGKε toward SAG. On average, phosphorylation of DDG in this fraction was about 5-fold lower than that of SAG in control cells. In contrast, DDG was efficiently phosphorylated by the cytosolic fraction harboring DGKs other than DGKε. This phosphorylation was not affected in DGKε-KD or DGKε-Myc-rescued cells (Fig. 2C, lower panel; and Fig. 2E). Taken together, the observed differences in the rate of the SAG-to-SAPA conversion between the tested cell variants reflected their different DGKε level. This indicates that the conversion of SAG to SAPA in these cells was catalyzed by DGKε.

### Depletion of DGKε abrogates TRIF-dependent cytokine production in LPS-stimulated cells

We next examined whether the DGKε depletion and re-introduction affected the LPS-induced responses, starting with the analysis of *Tlr4* and *Cd14* expression. The TLR4 and CD14 mRNA levels were similar in the DGKε-KD, DGKε-Myc-rescued, control, and maternal Raw264 cells, both before and after stimulation with LPS (100 ng/ml, 4 h). LPS induced an increase of CD14 mRNA abundance (1.4-1.9-fold, the highest in DGKε-KD cells) and a decrease of TLR4 mRNA (more than 2-fold) in all the cells (Fig. 3A, B).

**Figure 3.**
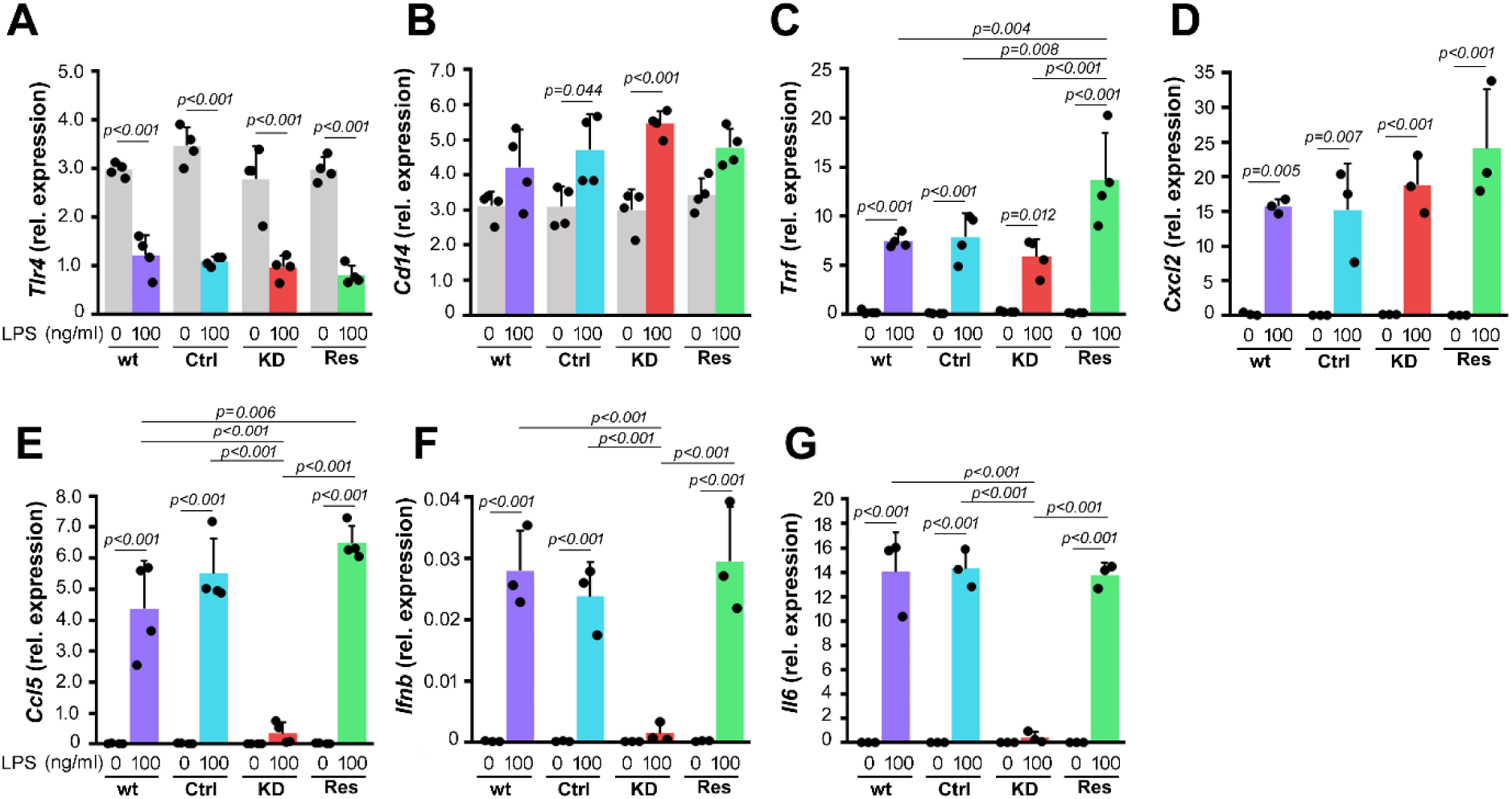
LPS-induced expression of TRIF-dependent cytokines is strongly down-regulated by *Dgke* knock-down and rescued by DGKε-Myc. Cells were left unstimulated or were stimulated with 100 ng/ml LPS (4 h, 37°C). (A-G) Transcripts of *Tlr4* (A), *Cd14* (B), *Tnf* (C), *Cxcl2* encoding MIP-2 (D), *Ccl5* encoding RANTES (E), *Ifnb* (F), and *Il-6* (G) were quantified by RT-qPCR relative to *Tbp* (A, D, F, G) or *Hprt* (B, C, E). Data shown are mean ± SD. Significantly different values as indicated by 1-way ANOVA with Tukey’s post hoc test are marked.

Despite the unaffected expression of *Tlr4* and *Cd14*, the LPS-stimulated production of cytokines was defective in cells depleted of DGKε, although to a various extent depending on the TLR4 signaling pathway involved. Thus, the mRNA level of TNFα, induced mainly in the MyD88-dependent manner, was reduced by about 25% (yet insignificantly) in comparison with wild-type Raw264 and Ctrl cells and was restored after DGKε re-introduction surpassing the relevant controls by 70-80% (Fig. 3C). The mRNA level of MIP-2, another cytokine produced in the MyD88-dependent pathway, was not affected (Fig. 3D). In contrast, the expression of genes encoding two strictly TRIF-dependent cytokines, CCL5/RANTES and IFN-β, and also IL-6 whose expression and mRNA stability are TRIF-dependent in macrophages (Kawai et al., 2001, Yamamoto et al., 2003, Bjorkbacka et al., 2004, Yanai et al., 2018, Nyati and Kishimoto 2022) were virtually nullified in the DGKε-KD cells. Their expression was fully restored in DGKε-Myc-rescued cells with that of *Ccl5* significantly surpassing maternal Raw264 cells by 48% (Fig. 3E-G). Overall, these results indicate that DGKε is required for efficient response of macrophages to LPS. Additionally, we confirmed that the obtained Ctrl and DGKε-Myc-rescued cells reasonably resembled the maternal Raw264 cells in their LPS-induced production of pro-inflammatory cytokines, with some increased potency observed in the rescued cells.

We pursued the studies by determining the secretion of TNFα and RANTES in cells stimulated with 10 and 100 ng/ml LPS for 4 h. The depletion of DGKε abrogated the TNFα production induced by 10 ng/ml LPS and the RANTES production at the both LPS concentrations (Fig. 4A, B). Notably, at 100 ng/ml LPS the TNFα production was only insignificantly lower in DGKε-KD cells (by about 39-46%) *vs.* Raw264 and Ctrl cells (Fig. 4A), in agreement with the TNFα mRNA assessments (Fig. 3C) Importantly, the re-introduction of DGKε restored the TNFα and RANTES production to levels insignificantly exceeding their production in Raw264 and Ctrl cells stimulated with 10 or 100 ng/ml LPS (Fig. 4A, B).

**Figure 4.**
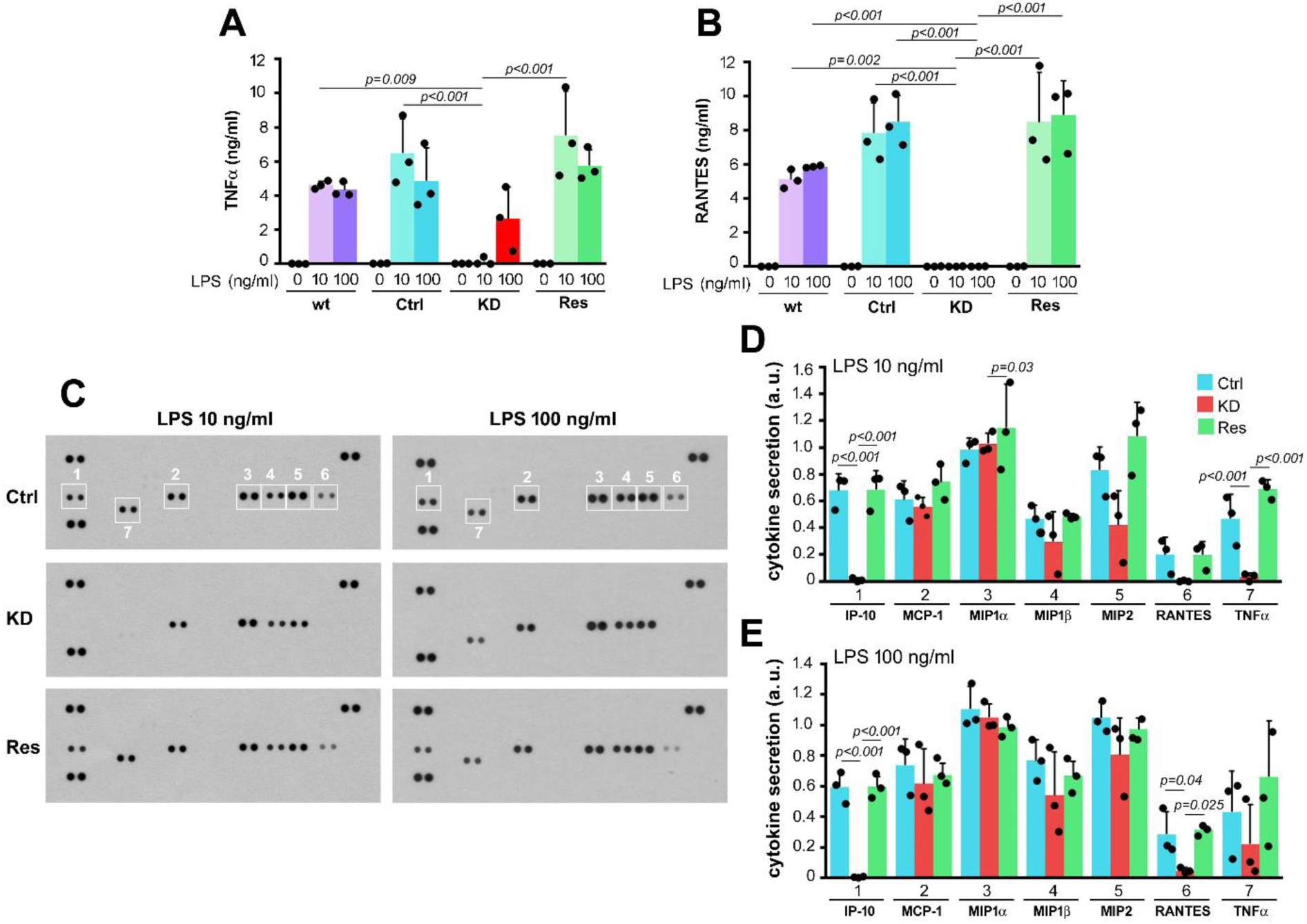
LPS-induced production of TRIF-dependent cytokines is dependent on DGKε. Cells were left unstimulated or were stimulated with 10 ng/ml or 100 ng/ml LPS (4 h, 37°C). (A, B) The concentration of TNFα (A) and CCL5/RANTES (B) in culture supernatants determined by ELISA. (C-E) Cytokine production in cells stimulated with 10 ng/ml or 100 ng/ml LPS determined using a cytokine array (C) and quantified by densitometric analysis of the array (D, E). Dots left unmarked in (C, upper panels) served as internal standards used to normalize signals between membranes. Data shown are mean ± SD. Significantly different values as indicated by 1-way ANOVA with Tukey’s post hoc test are marked. Additionally, in (A, B) all stimulated samples except for KD cells are significantly different from respective unstimulated ones (not marked).

These observations were confirmed by an analysis of the production of a whole array of inflammatory markers in cells stimulated with 10 and 100 ng/ml LPS. We found a virtual abrogation of the secretion of IP-10 and RANTES (both TRIF-dependent) in DGKε-KD cells and its full restoration in the DGKε-Myc-resued cells (Fig. 4C-E). In contrast, the production of the mainly MyD88-dependent MCP-1, MIP1α and β, MIP2, and TNFα tended to be inhibited slightly, reaching statistical significance for TNFα at 10 ng/ml LPS only (Fig. 4C-E). Taken together, the results indicated that the DGKε depletion abrogated the LPS-induced TRIF-dependent production of pro-inflammatory cytokines and to some extent inhibited also the MyD88-dependent one, especially at the lower LPS concentration.

### Depletion of DGKε inhibits CD14- and TRIF-dependent signaling of TLR4

To get a deeper insight into the contribution of DGKε to the MyD88- and TRIF-mediated signaling induced by TLR4, we followed the abundance of relevant proteins in the time course of the cell stimulation with LPS (1-4 h, 100 ng/ml LPS). Unexpectedly, we found a dramatic influence of DGKε on CD14 abundance; the mature forms of CD14 (bands estimated at 51-59 kDa which blurred in the presence of nonionic detergents, see Figs 7 and 8) with a slower gel migration caused by glycosylation (Stelter et al., 1996, Meng et al., 2008) were nearly absent in DGKε-KD cells, both resting and stimulated with LPS for up to 4 h. Instead, an accumulation of a faster-migrating doublet of CD14 (bands estimated at 47 and 51 kDa), tentatively assumed to represent immature forms of CD14, was found in these cells (Fig. 5A-C). Ctrl and DGKε-Myc-rescued cells differed slightly in the abundance of CD14 accumulation. Thus, in resting DGKε-Myc-rescued cells, mature CD14 was about 1.6-fold enriched relative to Ctrl cells, in agreement with the higher DGKε mRNA level and DGKε-mediated SAG phosphorylation in these cells (see Figs 1 and 2). The Ctrl cells responded to 4-hour LPS stimulation with a progressive accumulation of mature CD14. An increase, albeit insignificant, was also seen in DGKε-Myc-rescued cells for up to 2 h of stimulation (Fig. 5A, B). Thus, the depletion and reintroduction of DGKε affected the abundance of mature CD14 protein even though it did not modulate the *Cd14* gene expression. In contrast to CD14, TLR4 gradually disappeared in LPS-stimulated Ctrl and DGKε-Myc-rescued cells, owing to its CD14-dependent internalization and degradation (Husebye et al., 2006, Wang et al., 2007). Importantly, in the DGKε-KD cells lacking mature CD14, LPS stimulation did not induce the TLR4 disappearance (Fig. 5A, D).

**Figure 5.**
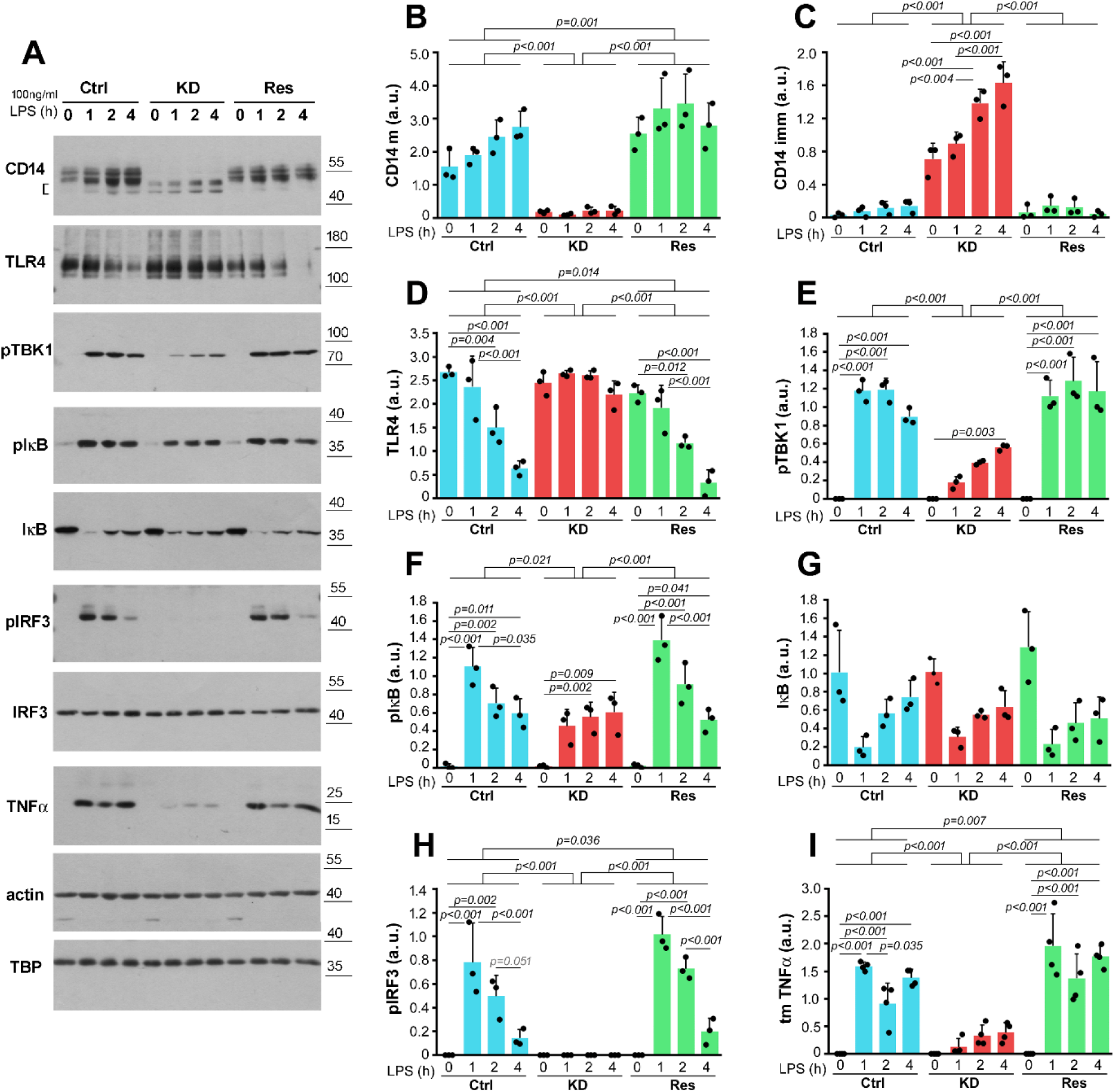
*Dgke* knock-down and rescue affect the abundance of mature CD14 and LPS-induced signaling. Cells were left unstimulated or were stimulated with 100 ng/ml LPS (1, 2 or 4 h, 37°C). (A) Abundance of indicated proteins in the cells determined by immunoblotting. Positions of molecular weight markers are shown on the right in kDa. Actin and TBP were visualized to verify equal loading of protein between wells. (B-I) Abundance of mature CD14 (CD14 m) (B), immature CD14 (CD14 imm, indicated by brackets) (C), TLR4 (D), phosphorylated TBK (pTBK) (E), phosphorylated IκB (pIκB) (F), IκB (G), phosphorylated IRF3 (pIRF3) (H), and transmembrane TNFα precursor (tm TNFα) (I) determined by densitometry. (B-D) and (F, G, I) analyzed relative to actin, (E) - to TBP, and (H) relative to IRF3. Data shown are mean ± SD. Significantly different values as indicated by two-way ANOVA with Tukey’s post hoc test are marked.

An analysis of the activity of NFκB and IRF3, two transcription factors reflecting, respectively, mainly the MyD88- and the strictly TRIF-dependent signaling pathways of TLR4 confirmed the limited dependence of the former and the strict dependence of the latter on the DGKε activity, and therefore on the abundance of mature CD14. The phosphorylation of IκB peaked at 1 h of LPS stimulation in both Ctrl and DGKε-Myc-rescued cells, and was lower by about 59-67% in DGKε-KD cells; its level was constant during the subsequent 3 h of stimulation in the DGKε-KD cells (Fig. 5F). IκB was degraded following its phosphorylation, with similar kinetics in all the cells (Fig. 5G). In turn, the phosphorylation of IRF3 was nullified in the DGKε-KD cells but was restored in the DGKε-Myc-rescued cells at a level moderately increased relative to Ctrl cells (Fig. 5K). Additionally, the activation (phosphorylation) of TBK1, a TRAF3 and TRAF6 effector (Everst et al., 2014, Liu et al., 2015, Tan and Kagan 2019), was significantly decreased in DGKε-KD cells relative to control throughout the 4-h period of LPS stimulation and fully restored in DGKε-Myc-rescued cells (Fig. 5E). A similar pattern was found for the production of TNFα transmembrane precursor (Fig. 5I).

Taken together, these results strongly support the relationship between DGKε abundance and activity, and the abundance of mature CD14 which ultimately determines the LPS-induced pro-inflammatory signaling.

### DGKε affects the surface level of CD14 and TLR4

To support further the above conclusion, we analyzed the level of cell-surface CD14 by flow cytometry. Only residual cell-surface CD14 colud be detected with an anti-CD14 antibody in DGKε-KD cells; CD14 re-appeared on the surface of DGKε-Myc-rescued cells at a level ca. 2-fold higher than in Ctrl cells, both before and after a 1-h stimulation with LPS (Fig. 6A). These findings were confirmed with the application of fluorescence-labeled inactive toxin aerolysin (FLAER) which binds selectively to the GPI anchor of a wide vairiety of GPI-linked proteins (Brodsky et al., 2000). The FLAER binding was reduced by about 87% in DGKε-KD compared with wild-type and Ctrl cells and was restored in DGKε-Myc-rescued cells to a level surpassing the control 1.7-fold. Notably, the labeling with FLAER was strongly decreased in both Ctrl and DGKε-Myc-rescued cells pretreated with PI-PLC (by about 78% and 69%, respectively), indicating that the detected protein(s) was indeed GPI-anchored in the plasma membrane (Fig. 6B).

**Figure 6.**
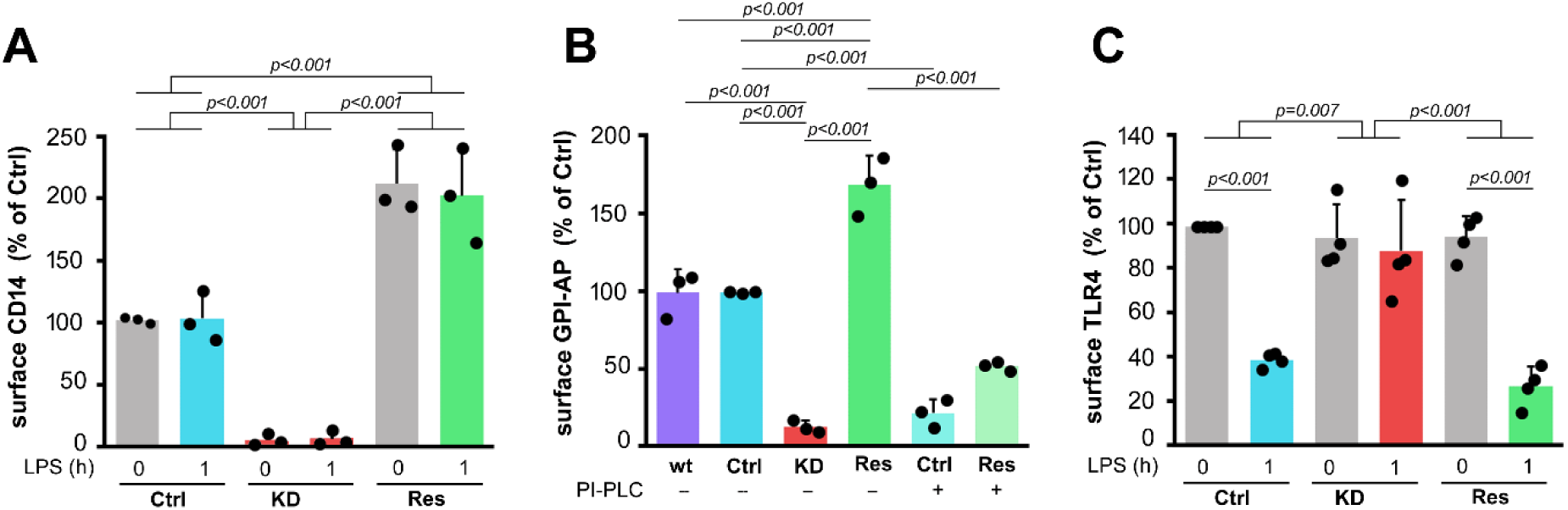
*Dgke* knock-down and rescue affect the cell surface level of CD14. Cells were left unstimulated or stimulated with 100 ng/ml LPS (1 h, 37°C). The cell-surface level of CD14 (A), total GPI-anchored proteins (GPI-AP) (B), and TLR4 (C) determined by flow cytometry. In (B), in a series of experiments unstimulated cells were pretreated with PI-PLC (0.2 U/ml, 1 h, 37°C) before labeling with FLAER. Data are expressed as a percentage of the value in unstimulated Ctrl cells. Significantly different values as indicated by two-way ANOVA (A, C) or one-way ANOVA (B) with Tukey’s post hoc test are marked.

In contrast to CD14, the surface level of TLR4 was comparable in all the cell types tested prior to LPS stimulation (Fig. 6C). The Ctrl and DGKε-Myc-rescued cells responded to LPS with a reduction of the cell surface level of TLR4 by 61% and 72%, respectively, while no change was found in DGKε-KD cells (Fig. 6C). These results corroborated the immunoblotting analysis (see Fig. 5) and confirmed the absence of TLR4 endocytosis upon LPS treatment in the DGKε-KD cells and its restitution following the re-introduction of DGKε.

### DGKε determines the production of GPI-anchored CD14

To get a further insight into the influence of DGKε on CD14 formation, we analyzed the distribution of CD14 in cell fractions obtained with the use of TX-100 and TX-114. The insolubility in cold TX-100 is a characteristic of raft proteins, including those with a GPI anchor (Plociennikowska et al., 2015). After fractionation of cells into TX-100-soluble, TX-100-insoluble (DRM), and SDS-soluble (cytoskeletal) fractions, the mature forms of CD14 (bands of ca. 51-59 kDa) were markedly enriched in the DRM fraction of maternal, Ctrl, and DGKε-Myc-rescued cells (Fig. 7A, C), as expected for a GPI-anchored protein (Matveichuk et al., 2024). Notably, in DGKε-KD cells CD14 was virtually absent in the DRM fraction (Fig. 7A, C). All the faster-migrating forms of CD14 in all the cell variants were detected exclusively in the TX-100-soluble fraction (Fig. 7A, B). This fraction also contained all of transferrin receptor and TLR4, as expected (Matveichuk et al., 2024), and DGKε-Myc in the case of the DGKε-Myc-rescued cells (Fig. 7A, D-F). For a reason currently unknown, the amount of transferrin receptor was reduced significantly in DGKε-Myc-rescued cells (Fig. 7F). Importantly, the overall abundance and enrichment in the DRM fraction of Lyn kinase was similar in all the cell lines (Fig. 7A, G), confirming the specificity of the effect of DGKε depletion toward the mature GPI-anchored CD14 form.

**Figure 7.**
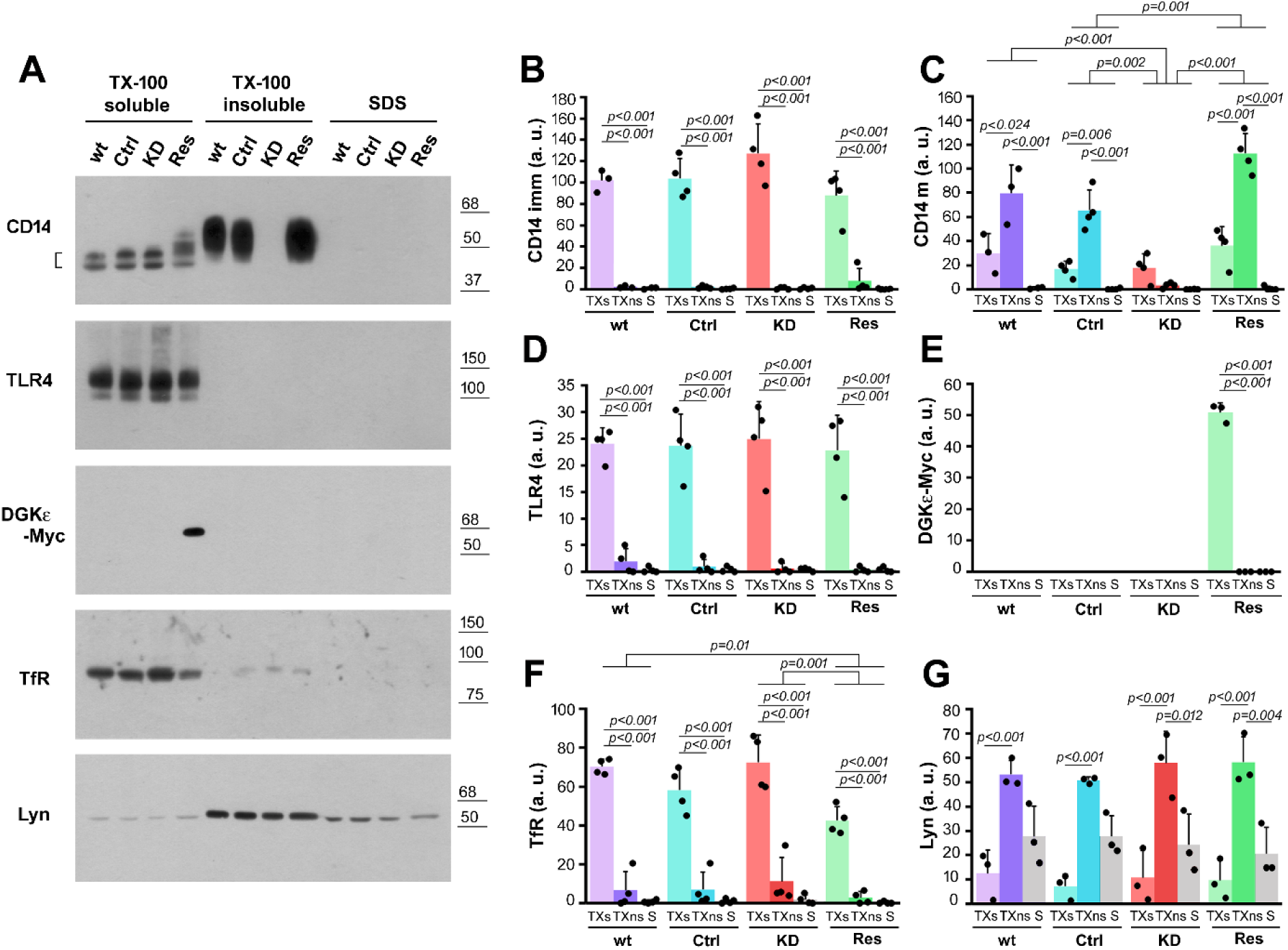
*Dgke* knock-down and rescue affect the abundance of mature forms of CD14. Cells were solubilized in 0.1% TX-100 and fractionated into TX-100 soluble (TXs), TX-100 insoluble (TXns) and SDS-soluble (S) fractions. (A) Distribution of indicated proteins in the fractions determined by immunoblotting. Positions of molecular weight markers are shown on the right in kDa. (B-I) Relative content of immature CD14 (CD14 imm, indicated by brackets) (B), mature CD14 (CD14 m) (C), TLR4 (D), DGKε-Myc (E), transferrin receptor (Tfr) (F) and Lyn (G) in each fraction determined by densitometry. Data shown are mean ± SD. Significantly different values as indicated by two-way ANOVA with Scheffe’s (C) or Tukey’s (F) post hoc test are marked. In (B, D, G) the two-way ANOVA indicated no significant differences between cell variants, therefore differences between fractions in a given cell variant were analyzed with one-way ANOVA with Scheffe’s (B) or Tukey’s (D, G) post hoc test. In (E) one-way ANOVA with Tukey’s post-hoc was used.

To pursue this subject further, we analyzed the distribution of CD14 in cell homogenate cytosolic and membrane fractions, with the latter subjected to solubilization and fractionation with TX-114, taking advantage of the partition of GPI-anchored proteins to the detergent fraction (Fujita et al., 2021). The faster-migrating doublet of CD14 (bands of about 47 and 51 kDa, corresponding to the TX-100-soluble forms of CD14 in Fig. 7) was found in the cytosolic fraction of all the cell variants (Fig. 8A). They were accompanied by small amounts of higher molecular weight forms (above 51 kDa) of CD14 likely corresponding to glycosylated soluble CD14 (Stelter et al., 1996). Only these soluble forms of CD14 were abundant in DGKε-KD cells (Fig. 8A). The cytosolic fraction accommodated all of IκB, as could be expected (Fig. 8A). To reveal the GPI-anchored CD14, the membrane-associated proteins were separated into hydrophilic and hydrophobic ones by solubilization with TX-114. The hydrophobic fraction contained the slow-migrating CD14 bands (ca. 51-59 kDa), present only in Ctrl and DGKε-Myc-rescued cells, which corresponded to the TX-100-insoluble ones (Fig. 8A, B, compare with Fig. 7). They were all shifted to aqueous phase after PI-PLC treatment cleaving the GPI moiety (Fig. 8A, B), confirming that these CD14 forms were anchored in membranes via GPI. Furthermore, considering the enzymatic activity of DGKε, their complete absence in DGKε-KD cells indicated a disturbed synthesis of GPI. Finally, the aqueous phase of all the cell types contained one dominant fast-migrating CD14 form (estimated at 47 kDa) (Fig. 8A). These would be a CD14 precursor containing the transmembrane moiety conferring its hydrophilic nature (Tanaka et al., 2004).

**Figure 8.**
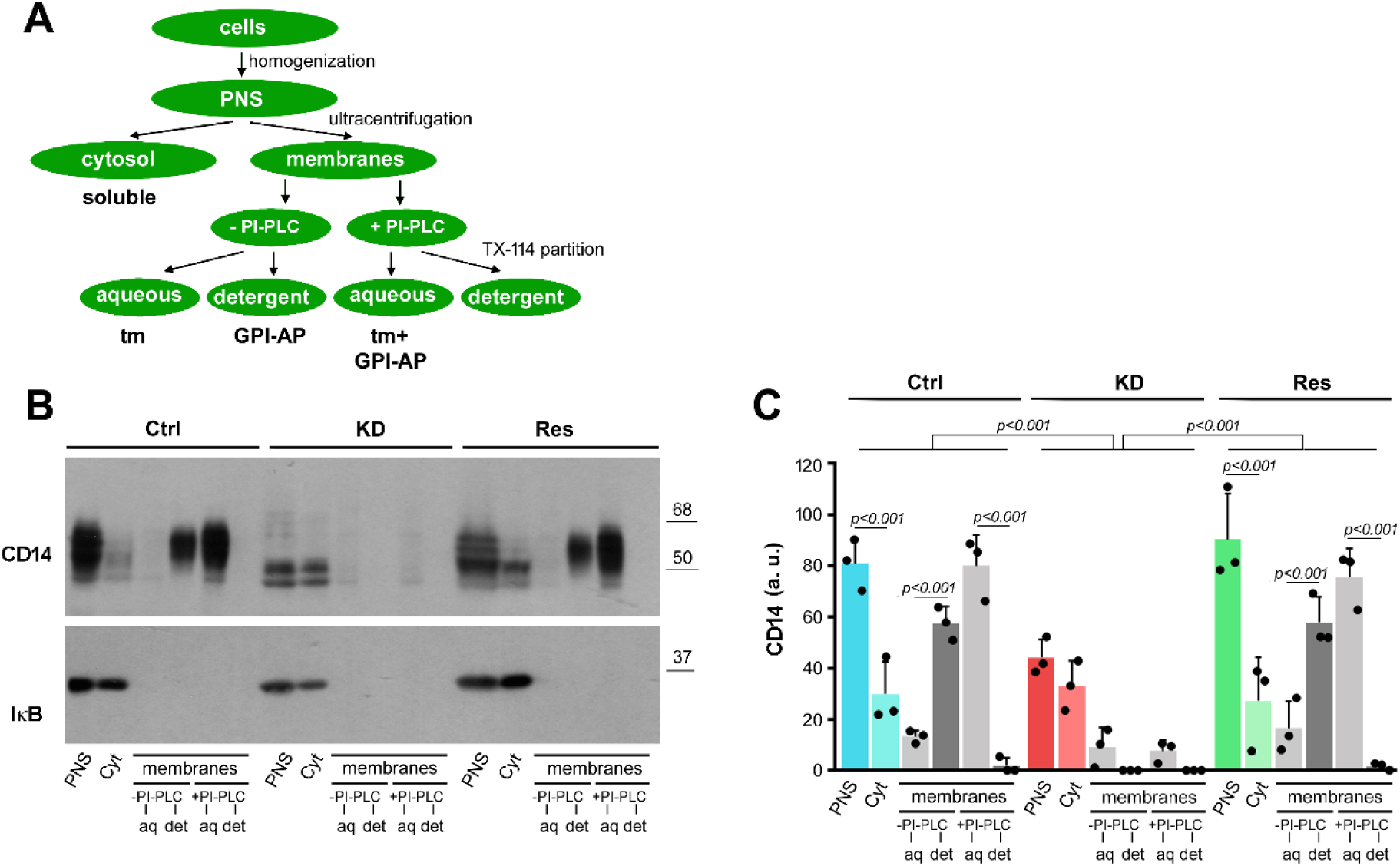
*Dgke* knock-down and rescue affect the abundance of GPI-anchored CD14. (A) Scheme of the cell fractionation. Post-nuclear supernatants (PNS) obtained from homogenates of Ctrl, DGKε-KD, and DGKε-Myc-rescued cells were fractionated into cytosolic (Cyt) and membrane fractions. The membrane fraction was treated or not with 2 U/ml PI-PLC, lysed in 2% TX-114 and partitioned into aqueous (aq) and detergent (det) phases. The fractions were subjected to SDS-PAGE in equivalent volumes. Localization of soluble, transmembrane (tm) and GPI-anchored CD14 (GPI-AP) is indicated. (B) Distribution of CD14 and IκB in cell fractions determined by immunoblotting. Positions of molecular weight markers are shown on the right in kDa. (C) Relative content of CD14 in cell fractions determined by densitometry. Data shown are mean ± SD. Significantly different values as indicated by two-way ANOVA with Tukey’s post hoc test are marked.

## DISCUSSION

CD14 is a GPI-anchored protein of the plasma membrane of myeloid cells, involved in TLR4 activation by facilitating LPS binding and governing endocytosis and endosomal signaling of TLR4 (Jiang et al., 2005, Zanoni et al., 2011). In the present study, we show that DGKε is required for the cell-surface presentation of GPI-linked CD14. A permanent depletion of Raw264 macrophage-like cells of DGKε led to the disappearance of mature CD14 equipped with a GPI anchor destined to the cell surface. Since a re-introduction of DGKε fully reversed this phenomenon, it is reasonable to conclude that it was indeed due to the deficiency of DGKε and not to some unspecified effect caused by *Dgke* silencing. The absence/reappearance of GPI-linked CD14 resulted in abrogation/recovery of the CD14-dependent pro-inflammatory signaling of TLR4, indicating DGKε as a key regulator of this signaling.

The DGKε depletion/rescue did not affect the mRNA levels of CD14 or TLR4 indicating that DGKε activity affected CD14 at a post-transcriptional stage. Specifically, the lack of DGKε led to the disappearance of the GPI-anchored mature forms of CD14 of a higher molecular weight due to glycosylation (Stelter et al., 1996), localized on the cell surface (Ciesielska et al., 2022). In control and DGKε-Myc-rescued cells, these CD14 forms were enriched in the TX-100-insoluble fraction of the cell lysates. They also partitioned into the detergent-rich phase of TX-114 after solubilizing membrane proteins, unless subjected to PI-PLC-mediated hydrolysis, which confirmed their membrane association via the GPI anchor. Taken together, these results indicate that in DGKε-depleted cells the maturation of CD14 was abrogated, likely because it failed to be modified with GPI. It has been established that for natively GPI-anchored proteins, the attachment of the GPI moiety is indispensable for their exit from the endoplasmic reticulum (Fujita and Kinoshita 2012), therefore, the anterograde transport of newly synthesized CD14 was probably prevented in DGKε-KD cells and followed partially by its degradation (Fujita and Kinoshita 2012) and partially by conversion into soluble CD14, as discussed below. This would ultimately prevent the replenishment of the cell surface pool of CD14 which is exhausted by the constitutive endocytosis and degradation of CD14 occurring in resting macrophages (Tan et al., 2015). Interestingly, in cells depleted of DGKε only the cytosolic doublet of anchor-less forms of CD14 was abundant. It resembled the doublet of truncated CD14 forms devoid of the C-terminal sequence, including the GPI attachment signal motif - the CD14-(1-335)-peptide described by (Stelter et al., 1996). These truncated CD14 forms underwent, however, complex *N*-glycosylation giving rise to higher-molecular weight forms of CD14 released from cells, thereby serving as a source of soluble CD14 detected in the cell milieu (Stelter et al., 1996). In view of our results, it seems likely that the anchor-less CD14 forms are produced by proteolysis of the transmembrane precursor of CD14 in the endoplasmic reticulum. In the absence of DGKε and the consequent GPI depletion, these are the only forms of CD14 synthesized by the cells. Interestingly, in patients suffering from paroxysmal nocturnal hemoglobinura (PNH) caused by a somatic mutation of the *PIGA* gene in hematopoietic stem cells which makes them defective in the GPI synthesis (see below), the serum level of soluble CD14 is normal (Schutt et al., 1995), which indicates that an alternative mechanism exists allowing its synthesis to compensates for the lack of shedding of mature GPI-linked CD14 from the cell surface (Bazil et al., 1991).

The biosynthesis of GPI is a multistep process initiated by the transfer of *N*-acetylglucosamine to the inositol ring of PI by a complex of phosphatidylinositol glycan anchor biosynthesis proteins (PIG proteins) with PIGA being the catalytic subunit. This first step of GPI formation, as well as subsequent de-*N*-acetylation of GlcNAc-PI to GlcN-PI, take place in the outer leaflet of the endoplasmic reticulum where PI is *de novo* synthesized from PA (Fujita and Kinoshita 2012, Kinoshita et al., 2020). The PA, in turn, is formed by two successive rounds of acylation of glycero-3-phosphate and lyso-PA. In the canonical pathway of PI synthesis, PA is transformed into CDP-DAG by cytidine diphosphate diacylglycerol synthase (CDS) 1 or 2, of which CDS2 prefers the SAPA. CDP-DAG is a short-lived intermediate used by phosphatidylinositol synthase (PIS) enzyme to form PI. Thus, this pathway seems to omit the SAG-to-SAPA phosphorylation (Bozelli and Epand 2019, Blunsom and Cockcroft 2020). It has recently been found, however, that the *de novo* PI synthesis is inhibited by the drug R59022 (Kim et al., 2022), which inhibits DGKα, DGKε, and DGKθ (Sato et al., 2013). This indicates that the *de novo* PI synthesis involves a cycle(s) of PA dephosphorylation and re-phosphorylation by the DGKs, which can be followed by acyl chain remodeling of PI via the Lands cycle to produce the C38:4 PI species, if the DAG phosphorylation is executed by DGK(s) other than DGKε (Kim et al., 2022). Our results show that the synthesis of GPI-anhored CD14 is tightly correlated with the presence and activity of DGKε only, indicating that the DGKε-mediated SAG-to-SAPA phosphorylation is an inevitable step in the synthesis of the PI pool from which the formation of GPI begins (Fig. 9).

**Figure 9.**
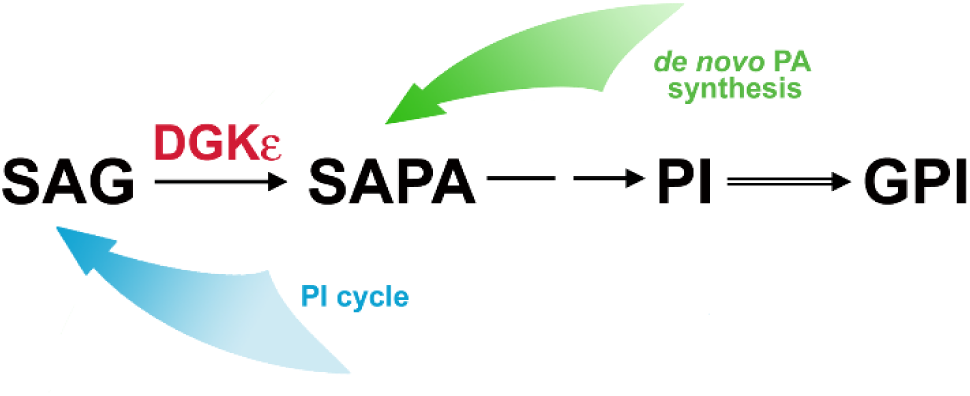
The SAG-to-SAPA phosphorylation by DGKε is required for PI synthesis, which is essential for GPI formation. The pathway can be supplied by intermediates derived from *de novo* PA synthesis and/or the PI cycle.

The results of Kim et al. (2022) obtained by metabolic labeling of HEK293 cells with radioactive PI precursors indicated that the *de novo* C38:4 PI synthesis, and also its re-synthesis following agonist-induced PI(4,5)P_2_ hydrolysis, occur with the contribution of multiple DGKs. In some cell types, the PI cycle was proposed to operate also under basal conditions, without agonist stimulation, utilizing pre-existing intermediates of the cycle (Barneda et al., 2022), potentially providing the PI required for the GPI production (Fig. 9). The two above-mentioned studies undermined the concept of an absolute requirement of the participation of DGKε in the PI synthesis, although the strict substrate specificity of DGKε predestines it to maintaining the C38:4 fatty acid signature of PI (Bozelli and Epand 2019). We addressed this issue by determining SAG phosphorylation in the membrane fraction of Raw264 cells depleted of peripheral DGKs with high salt extraction, and in the cytosolic fraction that in these conditions should accommodate all the DGKs except for DGKε. We confirmed the separation of DGKε from the other DGKs in the membrane fraction by a profound reduction of PA production from DAG lacking the C38:4 signature (DDG) and its presence in the cytosolic fraction. Notably, the knocking-down and restitution of DGKε affected the SAG-to-SAPA phosphorylation only in the membrane fraction, leaving that in the cytosol unaffected. The magnitude of the inhibition and recovery of the SAG phosphorylation in the membrane fraction closely followed the level of *Dgke* expression in the cell variants studied and was mirrored by the GPI-CD14 abundance. These results indicate that the PI synthesis involving the DGKε-mediated production of SAPA is crucial and likely non-redundant for the synthesis of the GPI anchor used for CD14 modification (Fig. 9). Recently, a sophisticated genetic screen has been performed in search of proteins involved in GPI biosynthesis. With the application of PIGA-KO HEK293 cells and chemically synthesized GlcNAc-PI, the CLPTM1L scramblase was found to aid the PIG-family proteins in the GPI biosynthetic pathway: CLPTM1L mediated the translocation of GlcN-PI to the luminal leaflet of the endoplasmic reticulum for further steps of the GPI biosynthesis (Wang et al., 2022). Our results place the DGKε activity upstream of PIGA and CLPTM1L in this pathway, which explains why it could not be discovered in the above-mentioned study of the Kinoshita’s group (2022).

When undertaking our study, we assumed that the depletion/rescue of DGKε expression would allow us to demonstrate its engagement in the biphasic generation of PI(4,5)P_2_ accompanying LPS binding to CD14 as part of a pro-inflammatory signaling cascade (Plociennikowska et al., 2016, Nguyen et al., 2013). As an example, the level of PI(4,5)P_2_ was decreased in the brain of DGKε-KO mice (Rodriguez de Turco et al., 2001). However, due to the unexpected effect of DGKε depletion on CD14 synthesis, its potential involvement in the PI cycle induced early (10-60 min) during stimulation of macrophages with 10-100 ng/ml LPS could not be determined directly.

The DGKε depletion and re-introduction affected the abundance of GPI-linked CD14, and consequently the induction of the TRIF-dependent endosomal signaling, which requires CD14-mediated endocytosis of TLR4 (Jiang et al., 2005, Zanoni et al., 2011). Also, the magnitude of MyD88-dependent signaling was affected, especially when a lower concentration of LPS was used for cell stimulation, reflecting the dispensability of CD14 involvement at high LPS doses (Gangloff et al., 2005, Borzecka et al., 2013). This line of data corroborates earlier findings showing that the pro-inflammatory signaling triggered by LPS is dependent on the abundance of plasma-membrane CD14. The cell-surface level of CD14 is determined by its complex cellular trafficking that differs substantially from that of TLR4 (Ciesielska et al., 2021), which can make CD14 sensitive to disturbances in this trafficking. Thus, we have recently found that the TLR4 signaling, especially the TRIF-dependent pathway, was reduced after inhibition of CD14 recycling or by up-regulation of the recycling followed by enhanced shedding of CD14, or by depletion of sphingomyelin (a raft lipid), all ultimately leading to the reduction of the total and cell-surface abundance of CD14 (Prymas et al., 2020, Ciesielska et al., 2022, Matveichuk et al., 2024). Furthermore, oxPAPC-mediated endocytosis of CD14 and its clearance from the cell surface inhibited the subsequent LPS-induced responses (Zanoni et al., 2017). However, the inhibitory effect of DGKε depletion on the abundance of GPI-CD14 was uniquely robust. Of utmost interest, the response of DGKε-KD and, consequently, CD14-depleted cells, to LPS mimicked the effect of the lack of PIGA in PNH patients, whose blood phagocytes displayed impaired responsiveness to low (1 ng/ml) but not high (100 ng/ml) concentrations of LPS (Duchow et al., 1993). It is worth noting that besides LPS, CD14 binds other ligands, including lipopeptides activating TLR2/TLR1 and TLR2/TLR6 (Borzecka et al., 2013, Matveichuk et al., 2024), and participates in the oxPAPC-dependent activation of the NLRP3 inflammasome (Zanoni et al., 2017) and non-canonical inflammasome by intracellular LPS (Vasudevan et al., 2022). The cell-surface CD14 also governs the endocytosis and pro-inflammatory activity of FAS receptors (Magri et al., 2024). These data suggest that the DGKε-dependent formation of GPI-linked CD14 likely affects several aspects of innate immunity.

Recently, an involvement of DGKε in energy lipid metabolism has been indicated by a series of studies conducted on DGKε-KO mice, in line with its localization in the endoplasmic reticulum. Thus, the DGKε-KO mice were prone to high-fat diet-induced obesity and insulin resistance due to alteration in triglyceride metabolizing enzymes in white adipocytes (Nakano et al., 2018); long-term high-fat diet feeding induced disturbances of brown adipose functioning (Nakano et al., 2020, 2024). In humans, recessive mutations of *DGKE,* some of which can lead to DGKε degradation (Traczyk et al., 2022), cause a kidney disease called atypical hemolytic uremic syndrome (aHUS) (Lemaire et al., 2013, 2021). Our results concerning the synthesis of GPI-anchored CD14 and LPS-induced pro-inflammatory signaling expand the list of DGKε-dependent processes even further.

## Supplemental data

This article contains supplemental data.

## Acknowledgments

The authors thank Dr. Jan Fronk (retired), formerly at the Faculty of Biology, University of Warsaw for helpful comments. We also thank Prof. Gianluca Baldanzi from the Center for Translational Research on Allergic and Autoimmune Diseases, University of Piemonte Orientale in Novara for NBD-labeled and unlabeled 1,2-didecanoyl-*sn*-glycerol used in studies on DGK activity.

## References

Aksoy E, Taboubi S, Torres D, Delbauve S, Hachani A, Whitehead MA, Pearce WP, Berenjeno IM, Nock G, Filloux A, Beyaert R, Flamand V, Vanhaesebroeck B. 2012. The p110δ isoform of the kinase PI(3)K controls the subcellular compartmentalization of TLR4 signaling and protects from endotoxic shock. Nature Immunology 13:1045–1054. DOI: 10.1038/ni.2426, PMID: 22922361

Balla T. 2013. Phosphoinositides: tiny lipids with giant impact on cell regulation. Physiological Reviews 93:1019–1137. DOI: 10.1152/physrev.00028.2012, PMID: 23899561

Barneda D, Janardan V, Niewczas I, Collins DM, Cosulich S, Clark J, Stephens LR, Hawkins PT. 2022. Acyl chain selection couples the consumption and synthesis of phosphoinositides. EMBO Journal 41:e110038. DOI: 10.15252/embj.2021110038, PMID: 35771169

Bazil V, Strominger JL. 1991. Shedding as a mechanism of down-modulation of CD14 on stimulated human monocytes. Journal of Immunol 147:1567–74. DOI: 10.4049/jimmunol.147.5.1567, PMID: 1880416

Bjorkbacka H, Fitzgerald KA, Huet F, Li X, Gregory JA, Lee MA, Ordija CM, Dowley NE, Golenbock DT, Freeman MW. 2004. The induction of macrophage gene expression by LPS predominantly utilizes Myd88-independent signaling cascades. Physiological Genomics 19:319–330. DOI: 10.1152/physiolgenomics.00128.2004, PMID: 15367729

Blunsom NJ, Cockcroft S. 2020. Phosphatidylinositol synthesis at the endoplasmic reticulum. *Biochimica et Biophysica Acta*, Molecular Cell Biology of Lipids 1865:158471. DOI: 10.1016/j.bbalip.2019.05.015, PMID: 31128095

Borzecka K, Plociennikowska A, Björkelund H, Sobota A, Kwiatkowska K. 2013. CD14 mediates binding of high doses of LPS but is dispensable for TNF-α production. Mediators of Inflammation 2013:824919. DOI: 10.1155/2013/824919, PMID 24453416

Bozelli JC Jr, Epand RM. 2019. Specificity of acyl chain composition of phosphatidylinositols. Proteomics 19:e1900138. DOI: 10.1002/pmic.201900138, PMID: 31207121

Brodsky RA, Mukhina GL, Li S, Nelson KL, Chiurazzi PL, Buckley JT, Borowitz MJ. 2000. Improved detection and characterization of paroxysmal nocturnal hemoglobinuria using fluorescent aerolysin. American Journal of Clinical Pathology 114:459–466. DOI: 10.1093/ajcp/114.3.459, PMID: 10989640

Cani PD, Bibiloni R, Knauf C, Waget A, Neyrinck AM, Delzenne NM, Burcelin R. 2008. Changes in gut microbiota control metabolic endotoxemia-induced inflammation in high-fat diet-induced obesity and diabetes in mice. Diabetes 57:1470–1481. DOI: 10.2337/db07-1403, PMID: 18305141

Chiang C-Y, Veckman V, Limmer K, David M. 2012. Phospholipase Cγ-2 and intracellular calcium are required for lipopolysaccharide-induced Toll-like receptor 4 (TLR4) endocytosis and interferon regulatory factor 3 (IRF3) activation. Journal of Biological Chemistry 287:3704–3709. DOI: 10.1074/jbc.C111.328559, PMID: 22194699

Ciesielska A, Krawczyk M, Sas-Nowosielska H, Hromada-Judycka A, Kwiatkowska K. 2022. CD14 recycling modulates LPS-induced inflammatory responses of murine macrophages. Traffic 23:310–330. DOI: 10.1111/tra.12842, PMID: 35503294

Ciesielska A, Matyjek M, Kwiatkowska K. 2021. TLR4 and CD14 trafficking and its influence on LPS-induced pro-inflammatory signaling. Cellular and Molecular Life Sciences 78:1233–1261. DOI: 10.1007/s00018-020-03656-y, PMID: 32914294

Cusson-Hermance N, Khurana S, Lee TH, Fitzgerald KA, Kelliher MA. 2005. Rip1 mediates the Trif-dependent Toll-like receptor 3- and 4-induced NF-κB activation but does not contribute to interferon regulatory factor 3 activation. Journal of Biological Chemistry 280:36560–36566. DOI: 10.1074/jbc.M506831200, PMID: 16118217

Dong R, Tan Y, Fan A, Liao Z, Liu H, Wei P. 2020. Molecular dynamics of the recruitment of immunoreceptor signaling module DAP12 homodimer to lipid raft boundary regulated by PIP_2_. Journal of Physical Chemistry B 124:504–510. DOI: 10.1021/acs.jpcb.9b11095, PMID: 31829096

Duchow J, Marchant A, Crusiaux A, Husson C, Alonso-Vega C, De Groote D, Neve P, Goldman M. 1993. Impaired phagocyte responses to lipopolysaccharide in paroxysmal nocturnal hemoglobinuria. Infection and Immunity 61:4280–4285. DOI: 10.1128/iai.61.10.4280-4285.1993, PMID: 8406889

Durieux JJ, Vita N, Popescu O, Guette F, Calzada-Wack J, Munker R, Schmidt RE, Lupker J, Ferrara P, Ziegler-Heitbrock HW, et al. 1994. The two soluble forms of the lipopolysaccharide receptor, CD14: characterization and release by normal human monocytes. European Journal of Immunology 24:2006–12. DOI: 10.1002/eji.1830240911, PMID: 8087168

Epand RM, So V, Jennings W, Khadka B, Gupta RS, Lemaire M 2016. Diacylglycerol kinase-ε: properties and biological roles. Frontiers Cell and Developmental Biology 4:112. DOI: 10.3389/fcell.2016.00112, PMID: 27803897

Everts B, Amiel E, Huang SC, Smith AM, Chang CH, Lam WY, Redmann V, Freitas TC, Blagih J, van der Windt GJ, Artyomov MN, Jones RG, Pearce EL, Pearce EJ. 2014. TLR-driven early glycolytic reprogramming via the kinases TBK1-IKKɛ supports the anabolic demands of dendritic cell activation. Nature Immunology 15:323–332. DOI: 10.1038/ni.2833, PMID: 24562310

Fitzgerald KA, Palsson-McDermott EM, Bowie AG, Jefferies CA, Mansell AS, Brady G, Brint E, Dunne A, Gray P, Harte MT, McMurray D, Smith DE, Sims JE, Bird TA, O’Neill LA. 2001. Mal (MyD88-adapter-like) is required for Toll-like receptor-4 signal transduction. Nature 413:78–83. DOI: 10.1038/35092578, PMID: 11544529

Fujita M, Kinoshita T. 2012. GPI-anchor remodeling: potential functions of GPI-anchors in intracellular trafficking and membrane dynamics. Biochimica et Biophysica Acta 1821:1050–1058. DOI: 10.1016/j.bbalip.2012.01.004, PMID: 22265715

Fujita M, Maeda Y, Kinoshita T. 2021. Partitioning of glycosylphosphatidylinositol (GPI)-anchored proteins with Triton X-114. In: Nishihara S, Angata K, Aoki-Kinoshita KF, Hirabayashi J, editors. Glycoscience Protocols (GlycoPODv2). Saitama (JP): Japan Consortium for Glycobiology and Glycotechnology. PMID: 37590577

Funda DP, Tucková L, Farré MA, Iwase T, Moro I, Tlaskalová-Hogenová H. 2001. CD14 is expressed and released as soluble CD14 by human intestinal epithelial cells in vitro: lipopolysaccharide activation of epithelial cells revisited. Infection and Immunity 69:3772–81. DOI: 10.1128/IAI.69.6.3772-3781.200, PMID: 11349042

Gangloff SC, Zähringer U, Blondin C, Guenounou M, Silver J, Goyert SM. 2005. Influence of CD14 on ligand interactions between lipopolysaccharide and its receptor complex. Journal of Immunology 175:3940–3945. DOI: 10.4049/jimmunol.175.6.3940, PMID: 16148141

Gil-de-Gómez L, Astudillo AM, Meana C, Rubio JM, Guijas C, Balboa MA, Balsinde J. 2013. A phosphatidylinositol species acutely generated by activated macrophages regulates innate immune responses. Journal of Immunology 190:5169–5177. DOI: 10.4049/jimmunol.1203494, PMID: 23585674

Gioannini TL, Zhang D, Teghanemt A, Weiss JP. 2002. An essential role for albumin in the interaction of endotoxin with lipopolysaccharide-binding protein and sCD14 and resultant cell activation. Journal of Biological Chemistry 277:47818–47825. DOI: 10.1074/jbc.M208786200, PMID: 12368295

Haziot A, Chen S, Ferrero E, Low MG, Silber R, Goyert SM. 1988. The monocyte differentiation antigen, CD14, is anchored to the cell membrane by a phosphatidylinositol linkage. Journal of Immunology 141:547–52. PMID: 3385210

Houjou T, Hayakawa J, Watanabe R, Tashima Y, Maeda Y, Kinoshita T, Taguchi R. 2007. Changes in molecular species profiles of glycosylphosphatidylinositol anchor precursors in early stages of biosynthesis. Journal of Lipid Research 48:1599–1606. DOI: 10.1194/jlr.M700095-JLR200, PMID: 17488725

Husebye H, Halaas R, Stenmark H, Tunheim G, Sandanger R, Bogen B, Brech A, Latz E, Espevik T. 2006. Endocytic pathways regulate Toll-like receptor 4 signaling and link innate and adaptive immunity. The EMBO Journal 25:683–692. DOI: 10.1038/sj.emboj.7600991, PMID: 16437160

Jiang Z, Georgel P, Du X, Shamel L, Sovath S, Mudd S, Huber M, Kalis C, Keck S, Galanos C, Freudenberg M, Beutler B. 2005. CD14 is required for MyD88-independent LPS signaling. Nature Immunology 6:565–570. DOI: 10.1038/ni1207, PMID: 15895089

Kagan JC, Medzhitov R. 2006. Phosphoinositide-mediated adaptor recruitment controls Toll-like receptor signaling. Cell 125:943–955. DOI: 10.1016/j.cell.2006.03.047, PMID: 16751101

Kagan JC, Su T, Horng T, Chow A, Akira S, Medzhitov R. 2008. TRAM couples endocytosis of Toll-like receptor 4 to the induction of interferon-beta. Nature Immunology 9:361–368. DOI: 10.1038/ni1569, PMID: 18345006

Kai M, Sakane F, Imai S, Wada I, Kanoh H. 1994. Molecular cloning of a diacylglycerol kinase isozyme predominantly expressed in human retina with a truncated and inactive enzyme expression in most other human cells. Journal of Biological Chemistry 269:18492–18498. DOI: 10.1016/S0021-9258(17)32484-0, PMID: 8027021

Kawai T, Adachi O, Ogawa T, Takeda K, Akira S. 1999. Unresponsiveness of MyD88-deficient mice to endotoxin. Immunity 11:115–122. DOI: 10.1016/S1074-7613(00)80086-2, PMID: 10435584

Kawai T, Ikegawa M, Ori D, Akira S. 2024. Decoding Toll-like receptors: Recent insights and perspectives in innate immunity. Immunity 57:649–673. DOI: 10.1016/j.immuni.2024.03.004, PMID: 37012345

Kawai T, Takeuchi O, Fujita T, Inoue J, Mühlradt PF, Sato S, Hoshino K, Akira S. 2001. Lipopolysaccharide stimulates the MyD88-independent pathway and results in activation of IFN-regulatory factor 3 and the expression of a subset of lipopolysaccharide-inducible genes. Journal of Immunology 167:5887–5894. DOI: 10.4049/jimmunol.167.10.5887, PMID: 11698465

Kelley SL, Lukk T, Nair SK, Tapping RI. 2013. The crystal structure of human soluble CD14 reveals a bent solenoid with a hydrophobic amino-terminal pocket. Journal of Immunology 190:1304–1311. DOI: 10.4049/jimmunol.1202446, PMID: 23264654

Kim J-I, Lee CJ, Jin MS, Lee C-H, Paik S-G, Lee H, Lee J-O. 2005. Crystal structure of CD14 and its implications for lipopolysaccharide signaling. Journal of Biological Chemistry 280:11347–11351. DOI: 10.1074/jbc.M414607200, PMID: 15632140

Kim YJ, Sengupta N, Sohn M, Mandal A, Pemberton JG, Choi U, Balla T. 2022 Metabolic routing maintains the unique fatty acid composition of phosphoinositides. EMBO Reports 23:54532. DOI: 10.15252/embr.202154532, PMID: 35712788

Kinoshita T. 2020. Biosynthesis and biology of mammalian GPI-anchored proteins. Open Biology 10:190290. DOI: 10.1098/rsob.190290, PMID: 32075561

Lee HC, Inoue T, Sasaki J, Kubo T, Matsuda S, Nakasaki Y, Hattori M, Tanaka F, Udagawa O, Kono N, Itoh T, Ogiso H, Taguchi R, Arita M, Sasaki T, Arai H. 2012. LPIAT1 regulates arachidonic acid content in phosphatidylinositol and is required for cortical lamination in mice. Molecular Biology of the Cell. 23:4689–700. DOI: 10.1091/mbc.E12-09-0673, PMID: 23097495

Lemaire M, Frémeaux-Bacchi V, Schaefer F, Choi M, Tang WH, Le Quintrec M, Fakhouri F, Taque S, Nobili F, Martinez F, et al. 2013. Recessive mutations in DGKE cause atypical hemolytic-uremic syndrome. Nature Genetics 45:531–536. DOI: 10.1038/ng.2590, PMID: 23542697

Lemaire M, Noone D, Lapeyraque AL, Licht C, Frémeaux-Bacchi V. 2021. Inherited Kidney Complement Diseases. Clinical Journal of the American Society of Nephrology. 16:942–956. doi: 10.2215/CJN.11830720, PMCID: PMC8216622

Li S, Huang F, Xia T, Shi Y, Yue T. 2023. Phosphatidylinositol 4,5-bisphosphate sensing lipid raft via inter-leaflet coupling regulated by acyl chain length of sphingomyelin. Langmuir 39:5995–6005. 10.1021/acs.langmuir.2c03492, PMID: 37086192

Liu S, Cai X, Wu J, Cong Q, Chen X, Li T, Du F, Ren J, Wu YT, Grishin NV, Chen ZJ. 2015. Phosphorylation of innate immune adaptor proteins MAVS, STING, and TRIF induces IRF3 activation. Science 347:aaa2630. DOI: 10.1126/science.aaa2630, PMID: 25636800

Lung M, Shulga, Ivanova YV, Ivanova PT, Myers DS, Milne S.B, Brown, HA, Tophma MK, Epand. RM 2009. Diacylglycerol kinase ε is selective for both acyl chains of phosphatidic acid or diacylglycerol. Journal of Biological Chemistry 284:31062–31073. DOI: 10.1074/jbc.M109.050617, PMID: 1974492

Magri Z, Jetton D, Muendlein HI, Connolly WM, Russell H, Smirnova I, Sharma S, Bunnell S, Poltorak A. 2024. CD14 is a decision-maker between Fas-mediated death and inflammation. Cell Reports 43:114685. DOI: 10.1016/j.celrep.2024.114685, PMID: 38843995

Mangan MSJ, Olhava EJ, Roush WR, Seidel HM, Glick GD, Latz E. 2018. Targeting the NLRP3 inflammasome in inflammatory diseases. Nature Reviews Drug Discovery 17:588–606. DOI: 10.1038/nrd.2018.97, PMID: 30006606

Matveichuk OV, Ciesielska A, Hromada-Judycka A, Nowak N, Ben Amor I, Traczyk G, Kwiatkowska K. 2024. Flotillins affect LPS-induced TLR4 signaling by modulating the trafficking and abundance of CD14. Cellular and Molecular Life Sciences 81:191. DOI: 10.1007/s00018-024-05221-3, PMID: 36804321

Meissner F, Scheltema RA, Mollenkopf H-J, Mann M. 2013. Direct proteomic quantification of the secretome of activated immune cells. Science 340:475–478. DOI: 10.1126/science.1232578, PMID: 23620012

Meng J, Parroche P, Golenbock DT, McKnight CJ. 2008. The differential impact of disulfide bonds and N-linked glycosylation on the stability and function of CD14. Journal of Biological Chemistry 283:3376–3384. DOI: 10.1074/jbc.M707640200, PMID: 18057036

Metz CN, Brunner G, Choi-Muira NH, Nguyen H, Gabrilove J, Caras IW, Altszuler N, Rifkin DB, Wilson EL, Davitz MA. 1994. Release of GPI-anchored membrane proteins by a cell-associated GPI-specific phospholipase D. The EMBO Journal 13:1741–51. DOI: 10.1002/j.1460-2075.1994.tb06438.x, PMID: 7512501

Motshwene PG, Moncrieffe MC, Grossmann JG, Kao C, Ayaluru M, Sandercock AM, Robinson CV, Latz E, Gay NJ. 2009. An oligomeric signaling platform formed by the Toll-like receptor signal transducers MyD88 and IRAK-4. Journal of Biological Chemistry 284:25404–11. DOI: 10.1074/jbc.M109.022392, PMID: 19605373

Nagaya H, Wada I, Jia YJ, Kanoh H. 2002. Diacylglycerol kinase delta suppresses ER-to-Golgi traffic via its SAM and PH domains. Molecular Biology of the Cell 13:302–316. DOI: 10.1091/mbc.01-05-0255, PMID:11809835

Nakano T, Seino K, Wakabayashi I, Stafforini DM, Topham MK, Goto K. 2018. Deletion of diacylglycerol kinase ε confers susceptibility to obesity via reduced lipolytic activity in murine adipocytes. FASEB Journal 32:4121–4131. DOI: 10.1096/fj.201701050R, PMID: 29509511

Nakano T, Suzuki A, Goto K. 2024. Ablation of diacylglycerol kinase ε promotes whitening of brown adipose tissue under high-fat diet feeding. Advances in Biological Regulation 91:100994. DOI: 10.1016/j.jbior.2023.100994, PMID: 37587945

Nakano T, Topham MK, Goto K. 2020. Mice lacking DGKε show increased beige adipogenesis in visceral white adipose tissue after long-term high-fat diet in a COX-2-dependent manner. Advances in Biological Regulation 75:100659. DOI: 10.1016/j.jbior.2019.100659, PMID: 31760328

Nguyen TT, Kim YM, Kim TD, Le OT, Kim JJ, Kang HC, Hasegawa H, Kanaho Y, Jou I, Lee SY. 2013. Phosphatidylinositol 4-phosphate 5-kinase α facilitates Toll-like receptor 4-mediated microglial inflammation through regulation of the Toll/interleukin-1 receptor domain-containing adaptor protein (TIRAP) location. Journal of Biological Chemistry 288:5645–5659. DOI: 10.1074/jbc.M112.410126, PMID: 23229526

Nyati KK, Kishimoto T. 2022. Recent advances in the role of Arid5a in immune diseases and cancer. Frontiers in Immunology 12:827611. DOI: 10.3389/fimmu.2021.827611, PMID: 35125794

Park BS, Song DH, Kim HM, Choi B-S, Lee H, Lee J-O. 2009. The structural basis of lipopolysaccharide recognition by the TLR4-MD-2 complex. Nature 458:1191–1195. DOI: 10.1038/nature07830, PMID: 19252480

Pettitt TR, Wakelam MJ. 1999 Diacylglycerol kinase ε, but not ζ, selectively removes polyunsaturated diacylglycerol, inducing altered protein kinase C distribution in vivo. Journal of Biological Chemistry 274:36181–36186. DOI: 10.1074/jbc.274.51.36181, PMID: 10593903

Plociennikowska A, Hromada-Judycka A, Borzecka K, Kwiatkowska K. 2015. Co-operation of TLR4 and raft proteins in LPS-induced pro-inflammatory signaling. Cellular and Molecular Life Sciences 72:557–581. DOI: 10.1007/s00018-014-1762-5, PMID: 25344887

Plociennikowska A, Hromada-Judycka A, Dembinska J, Roszczenko P, Ciesielska A, Kwiatkowska K. 2016. Contribution of CD14 and TLR4 to changes of the PI(4,5)P_2_ level in LPS-stimulated cells. Journal of Leukocyte Biology 100:1363–1373. DOI: 10.1189/jlb.2VMA1215-577R, PMID: 27534550

Posor Y, Jang W, Haucke V. 2022. Phosphoinositides as membrane organizers. Nature Reviews Molecular Cell Biology 23:797–816. DOI: 10.1038/s41580-022-00490-x, PMID: 35589852

Prymas K, Swiatkowska A, Traczyk G, Ziemlinska E, Dziewulska A, Ciesielska A, Kwiatkowska K. 2020. Sphingomyelin synthase activity affects TRIF-dependent signaling of Toll-like receptor 4 in cells stimulated with lipopolysaccharide. Biochimica et Biophysica Acta Molecular Cell Biology of Lipids 1865:158549. DOI: 10.1016/j.bbalip.2019.158549, PMID: 31730866

Rodriguez de Turco EB, Tang W, Topham MK, Sakane F, Marcheselli VL, Chen C, Taketomi A, Prescott SM, Bazan NG. 2001. Diacylglycerol kinase epsilon regulates seizure susceptibility and long-term potentiation through arachidonoyl-inositol lipid signaling. Proceedings of the National Academy of Sciences USA 98:4740–5. DOI: 10.1073/pnas.081536298, PMID: 11287665

Ryu J-K, Kim SJ, Rah S-H, Kang JI, Jung HE, Lee D, Lee HK, Lee J-O, Park BS, Yoon T-Y, Kim HM. 2017. Reconstruction of LPS transfer cascade reveals structural determinants within LBP, CD14, and TLR4-MD2 for efficient LPS recognition and transfer. Immunity 46:38–50. DOI: 10.1016/j.immuni.2016.11.007, PMID: 28061366

Sakane F, Murakami C, Sakai H. 2024. Upstream and downstream pathways of diacylglycerol kinase: Novel phosphatidylinositol turnover-independent signal transduction pathways. Advances in Biological Regulation 1:101054. DOI: 10.1016/j.jbior.2024.101054, PMID: 39368888

Sato M, Liu K, Sasaki S, Kunii N, Sakai H, Mizuno H, Saga H, Sakane F. 2013. Evaluations of the selectivities of the diacylglycerol kinase inhibitors R59022 and R59949 among diacylglycerol kinase isozymes using a new non-radioactive assay method. Pharmacology. 92:99–107. doi: 10.1159/000351849, PMID: 23949095

Sato S, Sugiyama M, Yamamoto M, Watanabe Y, Kawai T, Takeda K, Akira S. 2003. Toll/IL-1 receptor domain-containing adaptor inducing IFN-β (TRIF) associates with TNF receptor-associated factor 6 and TANK-binding kinase 1, and activates two distinct transcription factors, NF-κB and IFN-regulatory factor-3, in the Toll-like receptor signaling. Journal of Immunology 171:4304–4310. DOI: 10.4049/jimmunol.171.8.4304, PMID: 14530338

Shirai Y, Saito 2014. Diacylglycerol kinase as a possible therapeutic target for neuronal diseases. Journal of Biomedical Science 21:28. DOI: 10.1186/1423-0127-21-28, PMID: 24708409

Schutt C, Schilling T, Grunwald U, Stelter F, Witt S, Krüger C, Jack RS. 1995. Human monocytes lacking the membrane-bound form of the bacterial lipopolysaccharide (LPS) receptor CD14 can mount an LPS-induced oxidative burst response mediated by a soluble form of CD14. Research in Immunology 146:339–350. DOI: 10.1016/0923-2494(96)81038-8, PMID: 8719658

Simmons DL, Tan S, Tenen DG, Nicholson-Weller A, Seed B. 1989. Monocyte antigen CD14 is a phospholipid anchored membrane protein. Blood 73:284–9. PMID: 2462937

Sobocinska J, Roszczenko-Jasinska P, Zareba-Koziol M, Hromada-Judycka A, Matveichuk OV, Traczyk G, Lukasiuk K, Kwiatkowska K. 2018. Lipopolysaccharide upregulates palmitoylated enzymes of the phosphatidylinositol cycle: An insight from proteomic studies. Molecular and Cellular Proteomics 17:233–254. DOI: 10.1074/mcp.RA117.000050, PMID: 29118090

Stelter F, Pfister M, Bernheiden M, Jack RS, Bufler P, Engelmann H, Schutt C. 1996. The myeloid differentiation antigen CD14 is N- and O-glycosylated. European Journal of Biochemistry 236:457–464. DOI: 10.1111/j.1432-1033.1996.00457.x, PMID: 8617260

Su GL, Dorko K, Strom SC, Nüssler AK, Wang SC. 1999. CD14 expression and production by human hepatocytes. Journal of Hepatology 31:435–42. DOI: 10.1016/s0168-8278(99)80034-8, PMID: 10488701

Taguchi Y, Schätzl HM. 2014. Small-scale Triton X-114 extraction of hydrophobic proteins. Bio Protocol 4:e1139. DOI: 10.21769/BioProtoc.1139, PMID: 29170741

Tan Y, Kagan JC. 2019. Innate immune signaling organelles display natural and programmable signaling flexibility. Cell 177:384–398.e11. DOI: 10.1016/j.cell.2019.01.039, PMID: 30955886

Tan Y, Zanoni I, Cullen TW, Goodman AL, Kagan JC. 2015. Mechanisms of Toll-like Receptor 4 endocytosis reveal a common immune-evasion strategy used by pathogenic and commensal bacteria. Immunity 43:909–922. DOI: 10.1016/j.immuni.2015.10.008, PMID: 26588780

Tanaka S, Maeda Y, Tashima Y, Kinoshita T. 2004. Inositol deacylation of glycosylphosphatidylinositol-anchored proteins is mediated by mammalian PGAP1 and yeast Bst1p. Journal of Biological Chemistry 279:14256–63. DOI: 10.1074/jbc.M313755200, PMID: 14734546

Tang W, Bunting M, Zimmerman GA, McIntyre TM, Prescott SM. 1996. Molecular cloning of a novel human diacylglycerol kinase highly selective for arachidonate-containing substrates. Journal of Biological Chemistry 271:10237–10241. PMID: 8626589

Traczyk G, Hromada-Judycka A, Swiatkowska A, Wisniewska J, Ciesielska A, Kwiatkowska K. 2024. Diacylglycerol kinase-ε is *S*-palmitoylated on cysteine in the cytoplasmic end of its N-terminal transmembrane fragment. Journal of Lipid Research. 65:100480. DOI: 10.1016/j.jlr.2023.100480, PMID: 38008259

Traczyk G, Swiatkowska A, Hromada-Judycka A, Janikiewicz J, Kwiatkowska K. 2022. An intact zinc finger motif of the C1B domain is critical for stability and activity of diacylglycerol kinase-ε. International Journal of Biochemistry and Cell Biology 152:106295. DOI: 10.1016/j.biocel.2022.106295, PMID: 36356478

Traynor-Kaplan A, Kruse M, Dickson EJ, Dai G, Vivas O, Yu H, Whittington D, Hille B. 20177. Fatty-acyl chain profiles of cellular phosphoinositides. Biochimica et Biophysica Acta, Molecular and cell Biology of Lipids 1862:513–522. DOI: 10.1016/j.bbalip.2017.02.002, PMID: 28189644

Vasudevan SO, Russo AJ, Kumari P, Vanaja SK, Rathinam VA. 2022. A TLR4-independent critical role for CD14 in intracellular LPS sensing. Cell Reports 39:110755. DOI: 10.1016/j.celrep.2022.110755, PMID: 35508125

Violi F, Cammisotto V, Bartimoccia S, Pignatelli P, Carnevale R, Nocella C. 2023. Gut-derived low-grade endotoxemia, atherothrombosis and cardiovascular disease. Nature Reviews Cardiology 20:24–37. DOI: 10.1038/s41569-022-00737-2, PMID: 36216945

Virzi GM, Mattiotti M, de Cal M, Ronco C, Zanella M, De Rosa S. 2022. Endotoxin in sepsis: methods for LPS detection and the use of omics techniques. Diagnostics (Basel) 13:79. DOI: 10.3390/diagnostics13010079, PMID: 36673794

Wang Y, Chen T, Han C, He D, Liu H, An H, Cai Z, Cao X. 2007. Lysosome-associated small Rab GTPase Rab7b negatively regulates TLR4 signaling in macrophages by promoting lysosomal degradation of TLR4. Blood 110:962–971. DOI: 10.1182/blood-2007-01-066027, PMID: 17495131

Wang Y, Menon AK, Maki Y, Liu YS, Iwasaki Y, Fujita M, Guerrero PA, Silva DV, Seeberger PH, Murakami Y, Kinoshita T. 2022. Genome-wide CRISPR screen reveals CLPTM1L as a lipid scramblase required for efficient glycosylphosphatidylinositol biosynthesis. Proceedings of the National Academy of Sciences USA 119:e2115083119. DOI: 10.1073/pnas.2115083119, PMID: 35347902

Ware TB, Franks CE, Granade ME, Zhang M, Kim KB, Park KS, Gahlmann A, Harris TE, Hsu KL. 2020. Reprogramming fatty acyl specificity of lipid kinases via C1 domain engineering. Nature Chemical Biology 16:170–178. DOI: 10.1038/s41589-019-0445-9, PMID: 31932721

Yamamoto M, Sato S, Hemmi H, Uematsu S, Hoshino K, Kaisho T, Takeuchi O, Takeda K, Akira S. 2003. TRAM is specifically involved in the Toll-like receptor 4-mediated MyD88-independent signaling pathway. Nature Immunology 4:1144–1150. DOI: 10.1038/ni986, PMID: 14556004

Yamamoto M, Tanaka T, Hozumi Y, Saino-Saito S, Nakano T, Tajima K, Kato T, Goto K. 2014. Expression of mRNAs for the diacylglycerol kinase family in immune cells during an inflammatory reaction. Biomedical Research 35:61–68. DOI: 10.2220/biomedres.35.61, PMID: 24670677

Yanai H, Chiba S, Hangai S, Kometani K, Inoue A, Kimura Y, Abe T, Kiyonari H, Nishio J, Taguchi-Atarashi N, Mizushima Y, Negishi H, Grosschedl R, Taniguchi T. 2018. Revisiting the role of IRF3 in inflammation and immunity by conditional and specifically targeted gene ablation in mice. Proceedings of the National Academy of Sciences USA 115:5253–5258. DOI: 10.1073/pnas.1803936115, PMID: 29712861

Zanoni I, Ostuni R, Marek LR, Barresi S, Barbalat R, Barton GM, Granucci F, Kagan JC. 2011. CD14 controls the LPS-induced endocytosis of Toll-like receptor 4. Cell 147:868–880. DOI: 10.1016/j.cell.2011.09.051, PMID: 22078118

Zanoni I, Tan Y, Di Gioia M, Springstead JR, Kagan JC. 2017. By capturing inflammatory lipids released from dying cells, the receptor CD14 induces inflammasome-dependent phagocyte hyperactivation. Immunity 47:697–709.e3. DOI: 10.1016/j.immuni.2017.09.010, PMID: 29045901

