## Supplemental data for "Diacylglycerol kinase-ε is required for the formation of GPI-anchored CD14 and modulates the LPS-induced proinflammatory responses of macrophages"

**Supplemental Table 1. A**ntibodies used in the study

| Specificity of antibody | Host | Supplier | Catalog No. | Dilution | Application |
|---|---|---|---|---|---|
| Primary antibodies | | | | | |
| actin monoclonal IgG | Mouse | BD Biosciences | #612656 | 1:15000 | IB |
| CD14 monoclonal IgG | Rat | BD Pharmingem | #553738 | 1:4000-5000 | IB |
| calnexin | Goat | Abcam | #ab192439 | 1:1000 | IB |
| CD14 monoclonal IgG2a-FITC | Rat | eBioscience | #11-0141-82 | 1:200 | FC |
| IκBα monoclonal IgG | Mouse | Cell Signaling Techn. | #4814 | 1:1000 | IB |
| DGKζ | Rabbit | Abcam | #ab239081 | 1:1000 | IB |
| IRF3 monoclonal IgG | Rabbit | Cell Signaling Techn. | #4302 | 1:1000 | IB |
| Lyn | Rabbit | Cell Signaling Techn. | #2732 | 1:2000 | IB |
| Myc-tag monoclonal IgG | Mouse | Cell Signaling Techn. | #2276 | 1:1000 | IB |
| phospho-IκBα monoclonal IgG | Rabbit | Cell Signaling Techn. | #2859 | 1:1000 | IB |
| phospho-IRF3 monoclonal IgG | Rabbit | Cell Signaling Techn. | #4947 | 1:1000 | IB |
| phospho-TBK1/NAK monoclonal IgG | Rabbit | Cell Signaling Techn. | #5483 | 1:4000 | IB |
| TLR4 monoclonal IgG | Rabbit | Cell Signaling Techn. | #14358 | 1:1000 | IB |
| TLR4 monoclonal IgG2a-PE | Rat | BioLegend | #145404 | 1:200 | FC |
| TNFα monoclonal IgG | Rabbit | Cell Signaling Techn. | #11948 | 1:1000 | IB |
| transferrin receptor | Mouse | Invitrogen | #13-6800 | 1:2000-4000 | IB |
| Secondary antibodies | | | | | |
| mouse IgG-HRP | Goat | Jackson ImmunoResearch | # 115-035-003 | 1:6000-30000 | IB |
| rabbit IgG-HRP | Goat | Rockland | # 611-1302 | 1:8000 | IB |
| rat IgG-HRP | Goat | Merck | # A9037 | 1:10000 | IB |
| sheep IgG-HRP polyclonal IgG | Donkey | Jackson ImmunoResearch | # 713-035-003 | 1:6000 | IB |

IB, immunoblotting; FC, flow cytometry

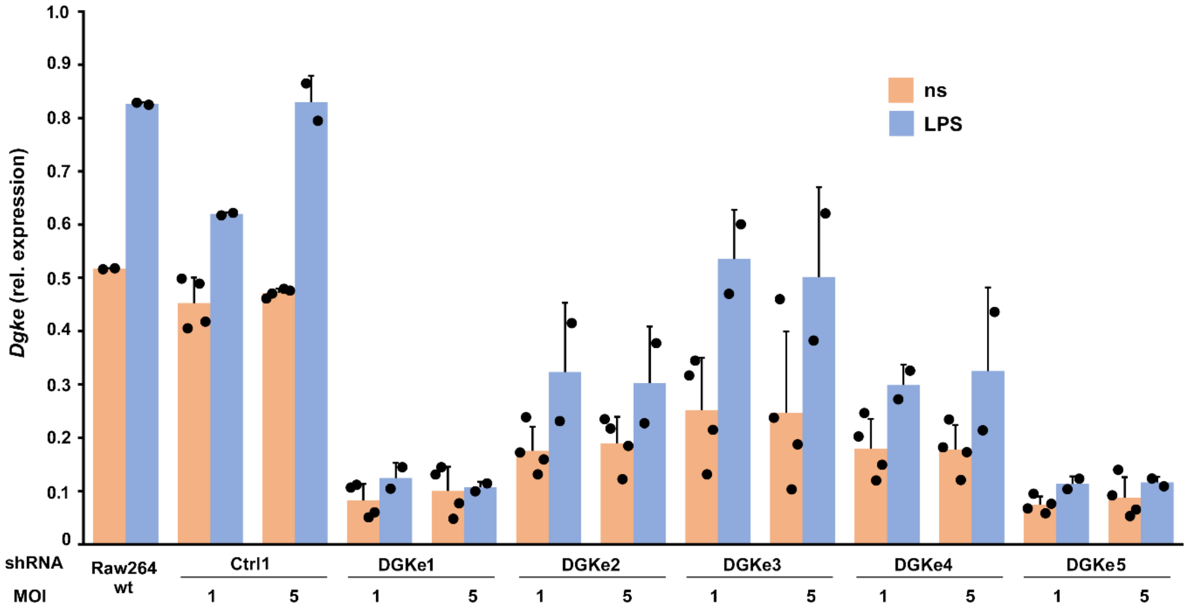


**Supplemental Fig. 1.** RT-qPCR analysis of DGKε mRNA in Raw264 cells after the knock-down of *Dgke* with shRNA. To silence *Dgke* expression, commercially available lentiviral particles bearing five shRNA variants were used individually (DGKe1-5) at MOI = 1 or 5, followed by puromycin selection. In parallel, control shRNA (Ctrl1) was applied at MOI = 1 or 5. Wild-type Raw264 cells – Raw264 wt. Cells were left unstimulated (ns) or were stimulated with 100 ng/ml LPS for 4 h. DGKε mRNA was quantified relative to TBP mRNA. Data shown are mean ± SD.

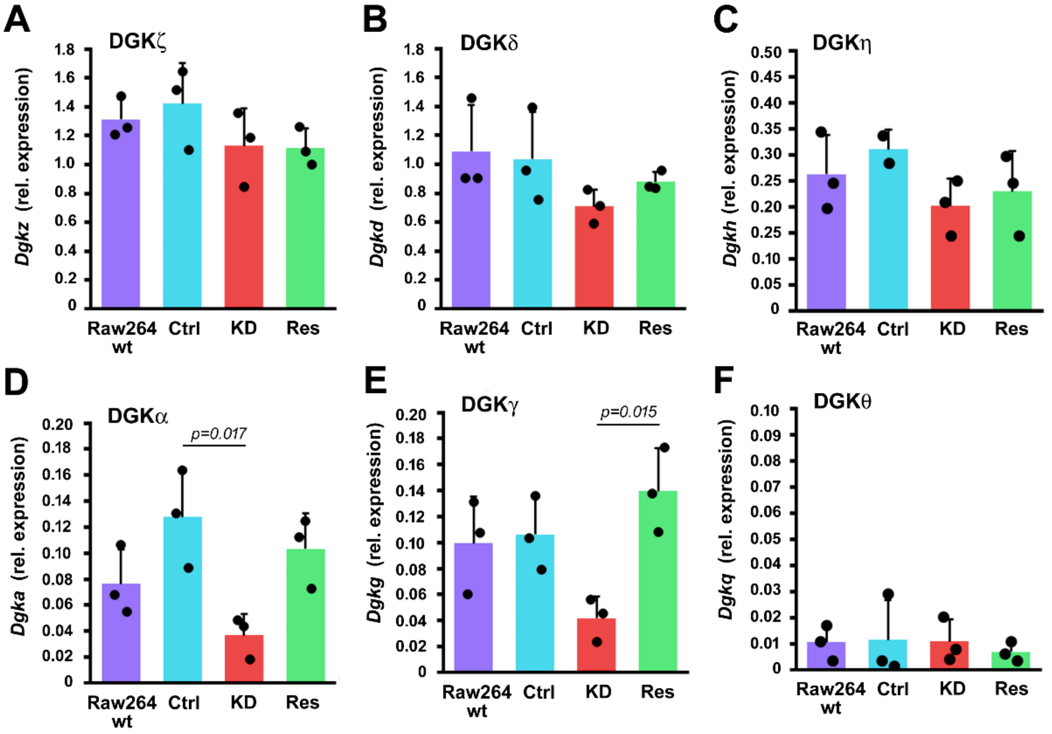


**Supplemental Fig. 2**. Expression of DGKs in Raw264 cells and their derivatives. Transcripts of *Dgkz* (A), *Dgkd* (B), *Dgkh* (C), *Dgka* (D), *Dgkg* (E), and *Dgkq* (F) were quantified relative to *Tbp*. Data shown are mean ± SD. Significantly different values as indicated by 1-way ANOVA with Tukey's post hoc test are marked.

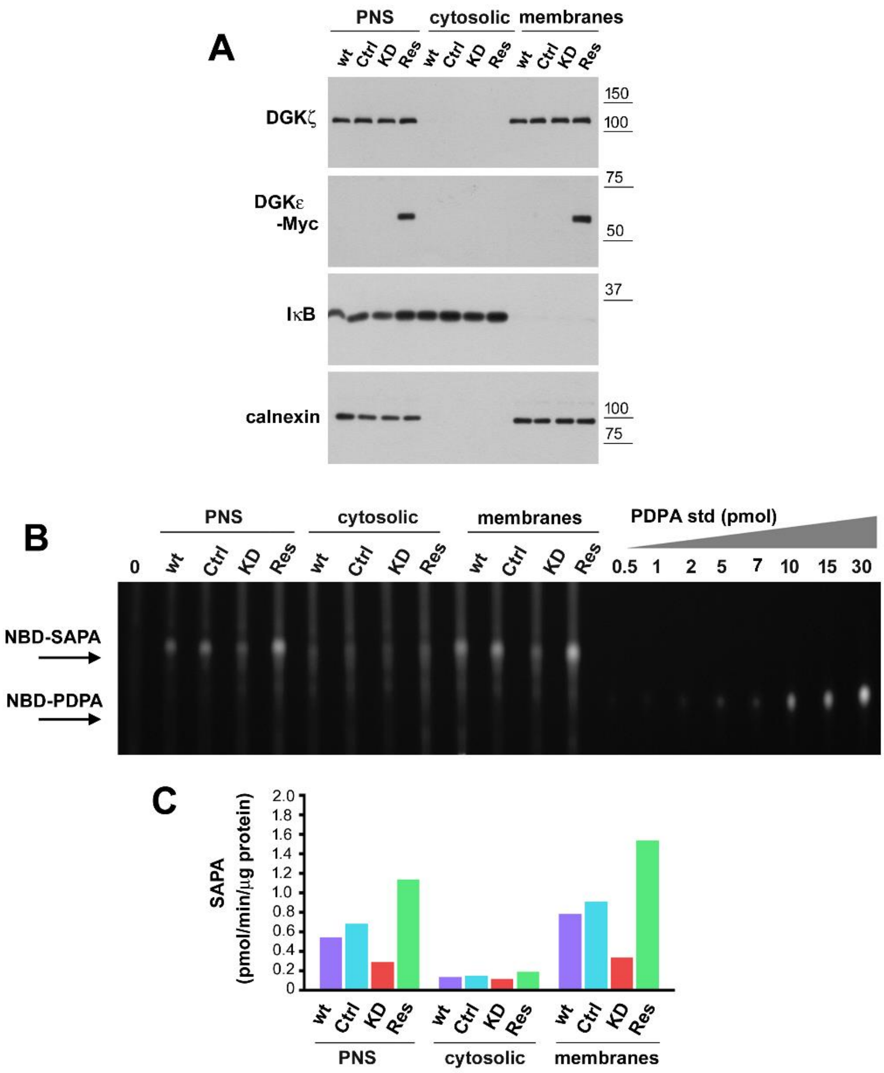


**Supplemental Fig. 3.** SAG phosphorylation in cell fractions not treated with 1 M NaCl. Post-nuclear supernatants (PNS) of wild-type Raw264 (wt), Ctrl, DGKε-KD, and DGKε-Myc-rescued cell homogenates were fractionated into cytosolic and membrane fractions without addition of 1 M NaCl. (A) Abundance of indicated proteins in cell fractions determined by immunoblotting. Positions of molecular weight markers are shown on the right in kDa. (B, C) Phosphorylation of SAG to SAPA in cell fractions determined using the fluorescence assay. (C) Representative TLC results revealing NBD-SAPA production. The reaction mixture contained 50 μg of total protein, lipids from 1/5 of the reaction mixture were applied onto the plate. NBD-labeled lipids were separated by TLC together with 0.5–30 pmol NBD-PDPA used to draw a standard curve for each experiment. “0” – no homogenate added. (C) SAPA production based on densitometric analysis of SAPA  and calibration curve for NBD-PDPA
